# THO and TRAMP complexes prevent transcription-replication conflicts, DNA breaks, and CAG repeat contractions

**DOI:** 10.1101/2021.12.06.471001

**Authors:** Rebecca E. Brown, Xiaofeng A. Su, Stacey Fair, Katherine Wu, Lauren Verra, Robyn Jong, Kristin Andrykovich, Catherine H. Freudenreich

**Author notes:** Contributed equally to this work (Rebecca E. Brown, Xiaofeng A. Su).

## Abstract

Expansion of structure-forming CAG/CTG repetitive sequences is the cause of several neurodegenerative disorders and deletion of repeats is a potential therapeutic strategy. Transcription-associated mechanisms are known to cause CAG repeat instability. In this study, we discovered that Thp2, an RNA export factor and member of the THO complex, and Trf4, a key component of the TRAMP complex involved in nuclear RNA degradation, are necessary to prevent CAG fragility and repeat contractions in a *S. cerevisiae* model system. Depletion of both Thp2 and Trf4 proteins causes a highly synergistic increase in CAG repeat fragility, indicating a complementary role of the THO and TRAMP complexes in preventing genome instability. Loss of either Thp2 or Trf4 causes an increase in RNA polymerase stalling at the CAG repeats and genome-wide transcription-replication conflicts (TRCs), implicating impairment of transcription elongation as a cause of CAG fragility and instability in their absence. Analysis of the effect of RNase H1 overexpression on CAG fragility and TRCs suggests that co-transcriptional R-loops are the main cause of CAG fragility in the *thp2Δ* mutants. In contrast, CAG fragility and TRCs in the *trf4Δ* mutant can be compensated for by RPA overexpression, suggesting that excess unprocessed RNA in TRAMP4 mutants leads to reduced RPA availability and high levels of TRCs. Our results show the importance of RNA surveillance pathways in preventing RNAPII stalling, TRCs, and DNA breaks, and show that RNA export and RNA decay factors work collaboratively to maintain genome stability.

## Introduction

Expansion-prone trinucleotide repeats (TNRs) are prone to DNA secondary structure formation, which can cause roadblocks to transcription or replication and interfere with DNA repair to cause repeat instability (changes in repeat length) and fragility (chromosome breaks) [1, 2]. Transcription through TNRs is an important *cis*-acting factor increasing repeat instability, and the known expansion-prone TNRs are transcribed within their associated genes [1, 3]. Tissue-specific RNA polymerase II (RNAPII) occupancy in the striatum and cerebellum during transcription elongation is highly associated with CAG repeat instability levels in Huntington’s disease (HD) mouse models [4, 5]. Transcription-coupled repair (TCR) has been reported to cause TNR instability in some systems [6]. However, TCR does not appear to be the major pathway causing expansions at the HD or Fragile X loci [1]. Therefore, other mechanisms of transcription-induced DNA damage and repair are likely also relevant.

Transcription through CAG and CGG repeats promotes R-loop formation and increases R-loop-dependent repeat instability [7-10]. R-loops promote single-stranded DNA (ssDNA) on the non-template strand, which is then available for secondary structure formation, and may provide a target for MutLγ nuclease cleavage and base excision repair [7, 11]. Therefore, formation of stable R-loops is one way in which transcription through G-rich repeats can cause chromosome fragility and instability. Another way that transcription can induce repeat instability is through changes in DNA supercoiling or chromatin structure that allow formation of secondary DNA structures. For example, remodeling by Isw1 during transcription is important in preventing CAG repeat expansions by helping to re-establish histone spacing after passage of RNAPII [12]. In addition, CAG or CTG slip-out structures can cause transcriptional arrest [13-15]. Despite the known importance of transcription in causing TNR instability, the impact of transcription-coupled RNA processing pathways on CAG repeat instability and fragility remains mostly unknown.

The yeast THO complex is a conserved eukaryotic transcription elongation factor which interacts with the nuclear pore-associated TREX complex to facilitate mRNA export [16]. THO is composed of four proteins, Tho2, Hpr1, Mft1, and Thp2, which form a highly stable complex, and Tex1, which is less tightly associated [17-19]. The THO complex travels co-transcriptionally with RNAPII to ensure stable messenger ribonucleoprotein (mRNP) formation and RNA extrusion during transcription [20, 21]. The THO-defective *hpr1Δ* mutant was shown to exhibit increased R-loop formation and transcription-associated recombination (TAR) [22]. Overexpression of RNase H1, the ribonuclease which directly degrades the RNA moiety in a RNA:DNA hybrid [23], abolishes the TAR phenotype in the *hpr1Δ* mutant [22]. THO mutants have a genome-wide accumulation of the Rrm3 protein, known to be required for replication through obstacles, in RNAPII transcribed genes [24-26]. The THO complex also counteracts telomere shortening and telomeric R-loop formation [27] and is needed for proper transcriptional elongation through the gene *FLO11*, that has internal tandem repeats [28]. Therefore, we hypothesized that the THO complex could play an important role in maintaining stability and reducing DNA breaks within CAG repeats by facilitating mRNA export and preventing R-loops to ensure normal transcription elongation.

The TRAMP (Trf4/5-Air1/2-Mtr4 Polyadenylation) complex is a functionally conserved nuclear RNA processing, degradation, and surveillance factor that facilitates degradation of RNA substrates by the addition of short unstructured poly(A) tails that target them for nuclear exosome-mediated degradation [29, 30]. TRAMP complexes are composed of the RNA helicase Mtr4, one of the non-canonical poly(A) polymerases Trf4 or Trf5 (forming TRAMP4 or TRAMP5 complexes respectively), and an RNA binding protein, either Air1 or Air2 [30]. Although TRAMP4 and TRAMP5 have similarity in their structures and RNA polyadenylation function, they also have specificity in RNA substrate species [30, 31]. The TRAMP complexes are important for degradation of rRNAs, tRNAs, small nuclear/small nucleolar (sn/sno) RNAs, and cryptic unstable transcripts (CUTs) by the exosome complex [31, 32]. *In vitro* studies have shown that the TRAMP adenylation and helicase activities act in a cooperative manner to unwind structured RNAs [30, 33]. The TRAMP complex has also been found to be co-transcriptionally recruited to promote rapid degradation of unwanted RNA transcripts, including spliced out introns, cryptic transcripts from rDNA regions, and aberrant mRNPs [32, 34, 35]. Deletion of yeast Trf4 causes nascent mRNA-mediated TAR, terminal deletions, and chromosome loss, which is suppressed by overexpression of RNase H1 [36, 37]. TAR is also evident in mutants lacking Rrp6, a major exoribonuclease of the nuclear exosome [33, 38], and *rrp6* mutants were shown to be associated with R-loop formation [39, 40]. Therefore emerging evidence has revealed new links between the TRAMP-exosome RNA decay machinery and genomic instability [41]. However, it is not known whether the TRAMP complexes are needed to maintain TNR stability. Also, although it is known that the TRAMP and THO complexes cooperate in controlling snoRNA expression [42], cross-talk between THO and TRAMP in regulating genetic instability has not been investigated.

In the course of screening for factors that protect against fragility of an expanded CAG tract in *Saccharomyces cerevisiae*, we identified members of both the THO and TRAMP complexes. We found that CAG repeat fragility and instability increase dramatically in the absence of either Thp2 or Mft1 of the THO complex or Trf4 or Rrp6 of the TRAMP/exosome machinery. Our data show that the THO and TRAMP complexes act in complementary pathways to reduce breakage at expanded CAG repeats in a manner distinct from accumulation of stable R-loops in the absence of RNase H1. Rather, defects in either complex result in RNAPII accumulation at the repeat tract and increased genome-wide transcription-replication conflicts (TRCs). Despite the similar phenotypes, genetic analysis indicates that the initial defect that leads to TRCs likely differs for THO and TRAMP mutants. In particular, defects in the TRAMP4 complex lead to reduced RPA availability, which is responsible for some of the TRCs and chromosome breaks. Our results highlight the importance of RNA export and processing factors in stabilizing expanded TNRs and preventing chromosome fragility at structure-forming repeats.

## Results

### The THO and TRAMP complexes both protect against chromosome breakage and repeat contractions

To study the role of RNA biogenesis and surveillance mechanisms on breakage and instability of an expanded CAG repeat tract, we used a previously established yeast artificial chromosome (YAC) system containing 70 CAG repeats (CAG-70) to evaluate the rate of fragility and frequency of repeat instability (Fig. 1A, [43]). When DNA double strand breaks (DSBs) occur at the CAG repeats, repair can occur by single-strand annealing or end joining to re-ligate the broken CAG repeat, causing contractions [44]. Alternatively, if repair fails or significant resection to the (G_4_T_4_)_13_ telomere seed occurs, this can result in *de novo* synthesis of a new telomere and YAC end loss (Fig. 1A). This event results in loss of the *URA3* gene, which confers resistance to 5-fluoroorotic acid (FOA). The rate of FOA resistance (FOA^R^), as a proxy for CAG repeat fragility, was measured by growing cells with a known starting repeat tract length for 6-7 generations without selection to allow fragility to occur, and then calculating the proportion of daughter yeast colonies that can grow on FOA media compared to non-selective media. Transcription occurs through the expanded CAG repeat tract on this YAC, with the majority of transcripts emanating from read-through transcription from the *URA3* gene (Fig. 1A), which generates a rCUG transcript [7, 12].

**Figure 1.**
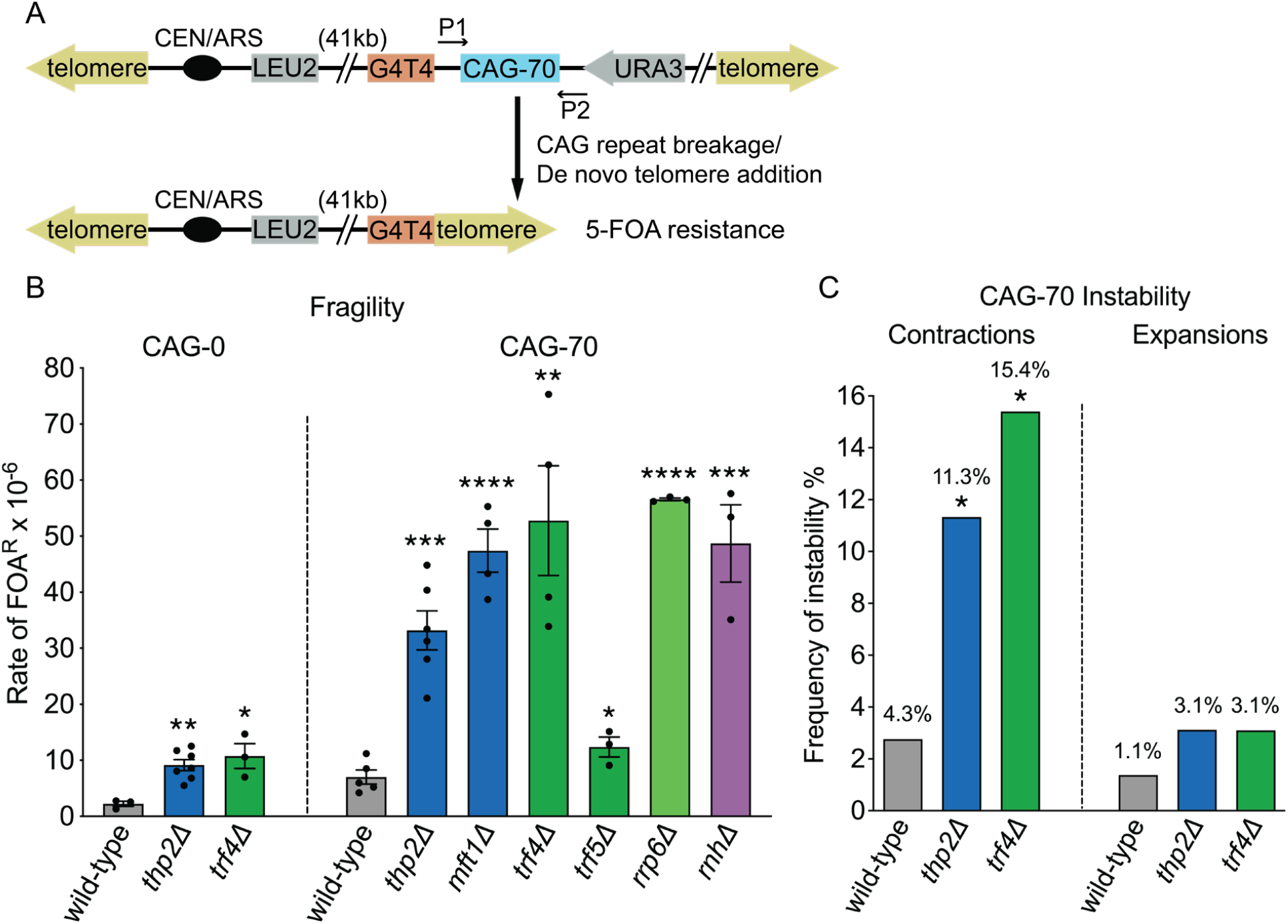
CAG repeat fragility increases in mutant strains defective for THO (blue bars), TRAMP4 and Rrp6 (green bars). (A) Assay system for CAG fragility. P1, P2 with arrows indicate the site-specific primers used for sizing CAG repeats by PCR amplification. (B) Rate of FOA^R^ × 10^− 6^ in indicated mutants; *Δ*, gene knock-out by replacing the target gene ORF with a selectable marker; each dot represents an individual data point; *p<0.05, **p<0.01 and ***p<0.001, ****p<0.0001 compared to wild-type with same CAG tract, by *t*-test. Average of at least 3 experiments ± SEM (standard error of the mean) is shown (Table S1 and S2). *rnhΔ*: *rnh1Δrnh201Δ*, data from [7]. (C) CAG repeat instability is increased in the *thp2Δ* or *trf4Δ* mutants. CAG-70 contraction and expansion frequencies in indicated mutants, a minimum of 130 reactions per mutant; *p<0.05, compared to wild-type, Fisher’s exact test (Table S3).

We initially identified a *thp2Δ* mutant as a top hit in a screen for yeast gene deletions that increase fragility of a CAG-85 repeat tract (see supplement of [45] for screen details). To confirm this phenotype, we deleted Thp2, a subunit of the THO complex, in a different strain background with a YAC carrying 70 CAG repeats and found that the *thp2Δ* strain exhibited a significant 4.7-fold increase in the rate of fragility compared to the wild-type strain (Fig. 1B; Table S1). A CAG-0 control also shows a significant 4-fold increase in fragility in *thp2Δ* compared to the wild-type strain (Fig. 1B; Table S2), indicating that the *thp2Δ* fragility phenotype is not unique to expanded triplet repeats but is exacerbated for this fragile site. Contraction frequency of CAG repeats in the *thp2Δ* mutant also increases significantly, 2.6-fold over wild-type (Fig. 1C; Table S3). To confirm that FOA-resistance was due to YAC end loss and not to other mechanisms such as point mutations occurring at the *URA3* marker gene, the structure of the YAC in multiple independently derived FOA resistant colonies was examined by Southern blot (method described in [43]). The results showed a 100% end-loss frequency with all the YAC structures consistent with *de novo* telomere addition at the G_4_T_4_ seed sequence (Table S4), indicating that loss of *URA3* is the cause of increased FOA resistance in the *thp2Δ* mutant. To confirm the importance of the THO complex in preventing CAG repeat breakage, another THO subunit, Mft1, was deleted. We chose Mft1 as it was identified in a second iteration of the screen for CAG fragility. The *mft1Δ* strain also exhibited a significant increase in CAG-70 repeat fragility that was similar to the level observed in the *thp2Δ* mutant (Fig. 1B; Table S1). Hpr1 and Tho2 deletions confer a strong growth defect [46], and were therefore not tested. The similar *thp2Δ* and *mft1Δ* phenotype confirmed that a defective THO complex causes an increase in fragility of expanded CAG repeats.

In our previous study, we found that transcription through the expanded CAG repeats on the YAC occurs bidirectionally by both cryptic transcription from the left and read-through transcription from the *URA3* gene to the right of the CAG tract [7], which could generate non-coding and possibly unstable rCAG and rCUG repeat transcripts. Such excess CAG repeat-containing RNA could be targeted by the TRAMP complex for degradation by the nuclear exosome. To investigate whether this RNA surveillance mechanism has an impact on CAG repeat fragility, we deleted Trf4, a non-canonical poly(A)-polymerase of the TRAMP4 complex, in the YAC-containing strains. In the *trf4Δ* mutant, the rate of CAG-70 repeat fragility is significantly and dramatically elevated 7.5-fold compared to the wild-type control (Fig 1B; Table S1), and the CAG repeat contraction frequency is also significantly increased by 3.6-fold compared to the wild-type (Fig. 1C; Table S3). Loss of *URA3* was confirmed by PCR (Table S4), indicating that FOA resistance is due to CAG fragility causing loss of the right arm of the YAC. A significant elevation was also seen in the *trf4Δ* no-tract control with a 4.9-fold increase over the wild-type control (Fig 1B; Table S2), indicating that the TRAMP complex also protects against chromosome fragility at non-repetitive DNA. Notably, we did not see such a dramatic increase in fragility of the CAG repeat when we knocked out the other TRAMP polyA-polymerase, Trf5. Although the increase in CAG-70 fragility in the *trf5Δ* mutant is significant compared to the wild-type control, it is only elevated by 1.8-fold compared to 7.5-fold for the *trf4Δ* mutant (Fig. 1B; Table S1). In previous studies, Trf4 and Trf5 were shown to have functionally distinct roles in polyadenylation of different RNA species with only partial redundancy [47]. We infer that most excess transcripts of expanded CAG repeats are polyadenylated and targeted for exosome degradation by the TRAMP4 complex containing Trf4.

TRAMP functions together with Rrp6, a 3′–5′ riboexonuclease component of the nuclear exosome, and Lrp1/Rrp47 which forms a heterodimer with Rrp6 and regulates its exonucleolytic activity [48]. Mutants in Trf4, Rrp6, and Lrp1 were all found to be strong positives in subsequent iterations of the CAG-85 fragility screen (Tufts University Molecular Genetics Project Lab course 2015, 2017). Deletion of Rrp6 led to a significantly elevated rate of CAG-70 repeat fragility compared to the wild-type control (Fig 1B; Table S1). Though the similar level of fragility for *trf4Δ* and *rrp6Δ* mutants suggests that the TRAMP complex may protect against CAG fragility through the same pathway as Rrp6-mediated exosome degradation of excess rCUG transcripts, the rate of CAG fragility in the *trf4Δrrp6Δ* double mutant is approximately additive compared to each single mutant (Fig. 2A). This suggests that CAG repeat fragility caused by the lack of Trf4 or Rrp6 arises from two semi-independent pathways. This conclusion is supported by decreased viability of the double compared to each single mutant (Fig. 2B), suggesting that DNA damage at the genomic level is also protected by both TRAMP4 and the exosome. We conclude that a functional TRAMP4-mediated RNA degradation mechanism protects expanded CAG repeats against fragility and deletions.

**Figure 2.**
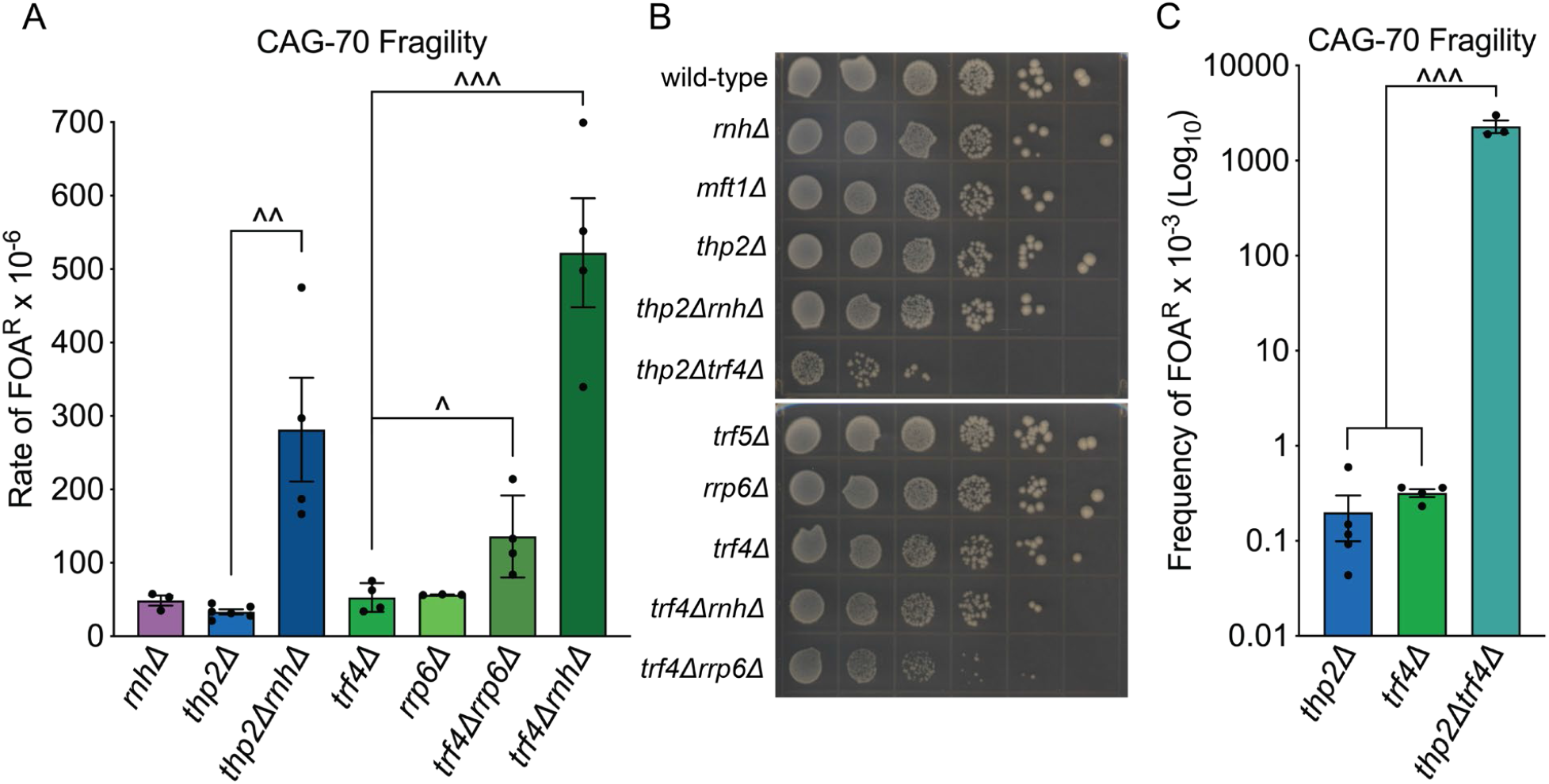
Loss of Thp2, Trf4, and RNase H synergistically increase CAG repeat fragility. (A) Rate of FOA^R^ × 10^−6^ in indicated mutants; *rnhΔ*: *rnh1Δrnh201Δ*, data from [7]; each dot represents an individual data point; ^p<0.05, ^^p<0.01, and ^^^p<0.001 compared to *thp2Δ* or *trf4Δ* single mutants, by *t*-test. Average of at least 3 experiments ± SEM is shown (Table S1). (B) Spot growth analysis. Indicated strains were plated on a selective synthetic media (YC-Leu-Ura) to maintain presence of the YAC. *rnhΔ*: *rnh1Δrnh201Δ*. The leftmost column contains the original plated cell suspension with OD_600_ of 1.0 for each individual strain with same volume (10 uL) plated, and serial dilution of 1:5 for the next column throughout the whole row. Plates were photographed 3 days post spotting. (C) Frequency of FOA^R^ × 10^−3^ in indicated mutants; each dot represents an individual data point; ^^^p<0.001: to *thp2Δ*: 1.1×10^−4^; to *trf4Δ*: 5.5×10^−4^, by *t*-test. Average of at least 3 experiments ± SEM is shown (Table S1).

In either *thp2Δ* or *trf4Δ* mutants, CAG repeat expansion frequency was only mildly increased in contrast to the significantly and dramatically elevated contraction frequency (Fig. 1C; Table S3). Therefore, there is a strong bias to contractions in THO and TRAMP mutants. Past research has shown that expansions often occur during gap repair pathways, whereas DSB repair within expanded CAG tracts most often lead to contractions [49, 50], therefore the bias to contractions is consistent with the high levels of breakage caused by deletion of Thp2 or Trf4.

### THO and TRAMP4 complexes cooperatively prevent fragility of expanded CAG repeats

In order to test if THO and TRAMP4 work in the same or different pathways in preventing CAG repeat fragility, we made a *thp2Δtrf4Δ* double mutant containing the CAG-70 YAC. These double mutants had a severe growth defect on normal yeast complete (YC) medium, compared to their isogenic wild-type control and *thp2Δ* or *trf4Δ* single mutants (Fig. 2B). The viability of *thp2Δtrf4Δ* double mutants in YC-Leu growth media to maintain the YAC is reduced to only 10%, almost an 8-fold decrease compared to the wild-type control (Fig. S1, Table S5). These results suggest that the THO and TRAMP complexes have complementary functions in processing RNA and that both are crucial for cell growth and fitness. Due to the low cell viability of the *thp2Δtrf4Δ* mutant, we were able to obtain only a few colonies carrying full-length CAG-70 tracts, therefore it was not feasible to perform our regular fragility protocol, which utilizes 10 single colonies per assay, with this double mutant. Therefore, we carried out a revised one-colony fragility assay as described in Methods and Materials. Similar control assays were also performed with the *thp2Δ* and *trf4Δ* single mutants. By using this modified fragility assay, we found that the frequency of FOA resistance drastically increases in the *thp2Δtrf4Δ* mutant, around 10,000-fold higher compared to the *thp2Δ* or *trf4Δ* single mutants (Fig. 2C; significance of p<0.0001 compared to either *thp2Δ* or *trf4Δ*). Such a high frequency of 5-FOA resistance in the double mutant provides evidence of a massive amount of DNA breakage at the repeats in the *thp2Δtrf4Δ* mutant and explains the low viability. The synergism between Thp2 and Trf4 in preventing CAG fragility indicates that THO and TRAMP4 act in complementary pathways to prevent breakage or repair breaks.

### R-loops play a partial role in causing CAG repeat fragility in the Thp2 mutant but no detectable role in a TRAMP4-defective background

In previous studies, abolishment of either the THO or TRAMP complex has been shown to increase R-loop-associated genome instability [22, 36, 37, 51]. Recently, the THO complex was shown to prevent R-loop formation throughout the cell cycle whereas other factors such as Senataxin only resolve R-loops during S-phase [52]. We previously showed that R-loops form preferentially at expanded CAG tracts *in vivo* and that increasing R-loop stability by removing both RNase H1 and RNase H2 proteins using a *rnh1Δrnh201Δ* double mutant further increased CAG repeat fragility [7]. Interestingly, the rate of CAG-70 repeat fragility in either *thp2Δ* or *trf4Δ* is very similar to the rate of the *rnh1Δrnh201Δ* (*rnhΔ*) mutant (Fig. 1B; Table S1). In order to test if CAG repeat fragility is caused by an R-loop-mediated mechanism in the *thp2Δ* or *trf4Δ* mutants, we created the *thp2Δrnh1Δrnh201Δ* and *trf4Δrnh1Δrnh201Δ* triple deletion strains. These two triple mutants do not show growth defects compared to the wild-type, *thp2Δ, trf4Δ*, or *rnhΔ* mutants (Fig. 2B). We found that CAG fragility in both *thp2ΔrnhΔ* and *trf4ΔrnhΔ* was significantly and synergistically elevated compared to their single mutants (Fig. 2A). These data demonstrate that accumulation of stable R-loops causes CAG repeat fragility through a different pathway than RNA biogenesis or degradation inefficiency due to defective THO or TRAMP4 complexes, respectively. However, we noted that the levels of CAG-70 fragility and synergism are different between *thp2ΔrnhΔ* and *trf4ΔrnhΔ*. The fragility for *thp2ΔrnhΔ* shows a 40-fold increase over the wild-type control and an 8.4-fold increase over *thp2Δ*, while the fragility for *trf4ΔrnhΔ* is even higher with a 74-fold increase over the wild-type control and a 10.5-fold increase over *trf4Δ* (Fig. 2A; Table S1). The somewhat lower level of CAG-70 fragility in the *thp2ΔrnhΔ* strain compared to the *trf4ΔrnhΔ* strain suggests that a portion of the CAG repeat fragility in the *thp2* mutant may be caused by R-loop formation or stabilization.

In order to test if R-loops physically accumulate at CAG repeats in either *thp2Δ* or *trf4Δ* mutants, we performed DNA:RNA immunoprecipitation coupled with qPCR (DRIP-qPCR) by using primers flanking the CAG repeats (Fig. 3A, Table S12; Methods and Materials). We were not able to detect a significant increase in R-loops at the CAG repeat in either *thp2Δ* or *trf4Δ* mutants (Fig. 3B, Table S5), unlike the 2-fold increase detected at the CAG-70 tract in a previous study for the *rnh1Δrnh201Δ* mutant [7]. We repeated the *rnhΔ* DRIP in parallel with the *thp2Δ* and *trf4Δ* DRIP to validate the efficacy of the S9.6 antibody and detected a 3.6-fold increase (Fig. 3B, Table S6), confirming the increased presence of R-loops in the *rnhΔ* strain and negative result for the *thp2Δ* and *trf4Δ* strains.

**Figure 3.**
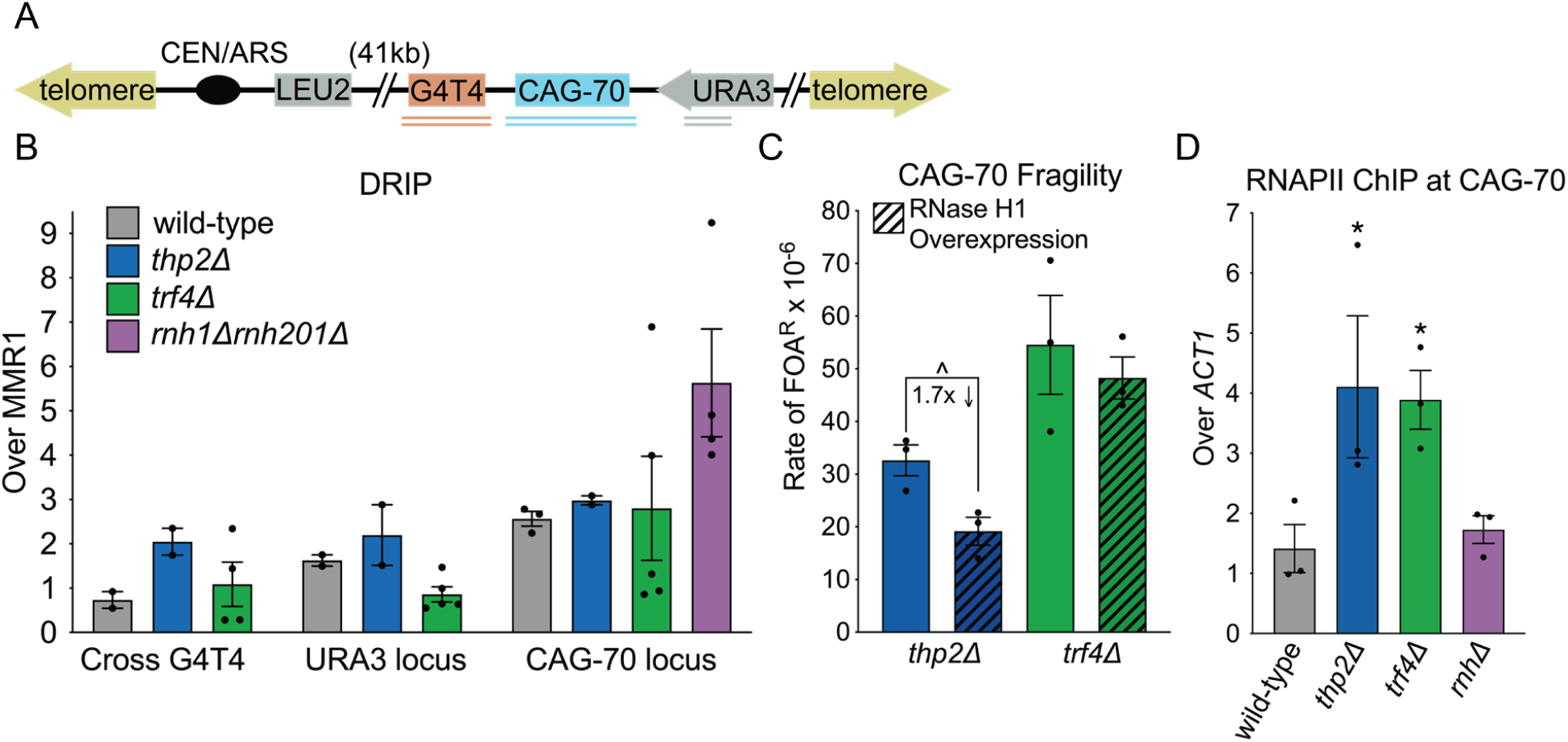
Analysis of R-loop presence and RNA polymerase stalling and effect on CAG-70 fragility in RNA biogenesis mutants. (A) YAC used for fragility and DNA:RNA immunoprecipitation (DRIP) assays. Double underlines indicate the qPCR amplicons used. (B) DRIP in wild-type, *thp2Δ, trf4Δ*, and *rnh1Δrnh201Δ* strains. Each bar represents the mean ± SEM of at least two experiments from two biological replicates; each dot represents an individual data point; *rnhΔ* mutant data is from [7] and one additional replicate used as a parallel control for S9.6 antibody (Table S6). (C) Rate of FOA^R^ × 10^−6^ in indicated mutants and RNase H1 overexpression conditions; each dot represents an individual data point; ^p<0.05, compared to no induction condition in the same mutant, by *t*-test. Average of at least 3 experiments ± SEM is shown (Table S1). See Fig. S2A for RNase H1 expression levels in overexpression strains (Table S7). (D) RNAPII chromatin immunoprecipitation (ChIP) at the CAG-70 repeat in the indicated strains, shown as the IP/INPUT signal of the RNAPII ChIP at the CAG repeat normalized to the IP/INPUT signal at the *ACT1* genomic locus. Each bar represents the mean ± SEM of at least three experiments (biological replicates); each dot represents an individual experiment; *p<0.05, compared to wild-type, by *t*-test (Table S8).

To test if R-loop formation influences CAG repeat fragility in the RNA biogenesis mutants using a functional assay, we inserted an inducible *MET25* promoter [53] before the *RNH1* gene, which encodes RNase H1, to allow induced overexpression of *RNH1* in synthetic medium lacking methionine. We confirmed overexpression of RNase H1 by using a reverse-transcription reaction and quantitative PCR (qRT-PCR) in the wild-type background as well as *thp2Δ* and *trf4Δ* mutants (Fig S2A, Table S7). Fragility analysis showed a 40% reduction (1.7-fold decrease) in the rate of FOA resistance compared to the un-induced condition for the *thp2Δ* mutant (p=0.027; Fig. 3C, Table S1). These results indicate that depletion of Thp2 causes a functional accumulation R-loops at the CAG repeats, consistent with previous findings that R-loops accumulate in THO mutants [22, 27, 51, 54]. We conclude that R-loops within the CAG tract in the THO-defective mutant may be transient or of a short length since they were not detected by DRIP but do contribute to causing the increase in CAG fragility observed in this background. Castillo-Guzman and Chédin recently defined two classes of R-loops, promoter-paused (Class I) and elongation-associated (Class II). The types of R-loops that accumulate at the CAG repeat in the absence of Thp2 may be similar to Class I R-loops that are shorter, less stable, and not readily detected by DRIP [55].

In contrast, no reduction in the FOA^R^ rate was seen in the *trf4Δ* background upon RNase H1 over-expression (Fig 3C, Table S1). Combined with the failure to detect increased R-loops at the CAG tract by DRIP in the *trf4Δ* mutant, these results suggest that the increased CAG fragility in this background is not a result of R-loop accumulation but rather due to another mechanism.

### Loss of either THO or TRAMP4 results in RNA polymerase II accumulation at expanded CAG repeats and increased transcription-replication conflicts (TRCs) genome-wide

The largest RNAPII-subunit Rpb1 is targeted for degradation in the absence of THO subunit Tho2, suggesting RNAPII stalling occurs in THO-defective mutants [56]. To explore whether RNAPII was stalled at expanded CAG repeats in either *thp2Δ* or *trf4Δ* mutants, we performed RNAPII chromatin immunoprecipitation (ChIP) analysis to detect elongating RNAPII. Indeed, we discovered that RNAPII was significantly enriched at the CAG repeat tract compared to a control locus by about 3-fold in both the *thp2Δ* and *trf4Δ* mutants (Fig. 3D; p=0.05 *thp2Δ* to wild-type, p=0.017 *trf4Δ* to wild-type, Table S8). In contrast, we did not detect a significant increase in RNAPII enrichment at CAG-70 repeats in the *rnh1Δrnh201Δ* mutant (Fig. 3D). These results indicate that defective THO and TRAMP complexes cause RNAPII stalling and impaired transcription elongation at expanded CAG repeats.

One possible explanation for DSBs at the expanded CAG tract is TRCs due to a stalled RNA polymerase II complex (RNAPII) interfering with DNA replication [57]. To determine whether TRCs were occurring in *thp2Δ, trf4Δ*, or *rnh1Δrnh201Δ* mutants, we employed a proximity ligation assay (PLA) using antibodies against PCNA to detect the replisome, and RNAPII to detect the transcription machinery. This assay detects proteins that are within 40 nm of each other [58], which are detectable as foci in the nucleus (example nuclei are shown in Fig. 4A). The size of the foci is comparable to what has been observed for this assay in mammalian nuclei (see scale bars in Fig. 4A and in [59, 60] as examples). Quantification of the number of foci in each strain showed a highly significant increase between wild-type and each indicated mutant, with the effect being smallest for the *rnh1Δrnh201Δ* strain (1.3-fold increase, p<0.001), followed by a further significant 1.6-fold increase in the *thp2Δ* nuclei, and an even more striking 2.2-fold increase in *trf4Δ* nuclei (p<0.0001 for both) (Fig. 4B, Table S9). Some cells had foci detected outside of the nucleus, but the percentage of these compared to total foci counted within the nucleus was small and similar across strains (Table S10). These data indicate that all three mutants cause TRCs genome-wide, however the effect is significantly stronger in the THO and TRAMP mutants compared to the RNase H mutant strain. Interestingly, the PLA data correlate more closely with the levels of RNAPII detected by ChIP within the CAG-70 tract, and less well with the levels of R-loops within the repeat tract detected by DRIP. These data suggest that a stalled RNAPII, with or without an accompanying R-loop, is a stronger barrier to DNA replication than R-loops that persist in the absence of RNase H processing.

**Figure 4.**
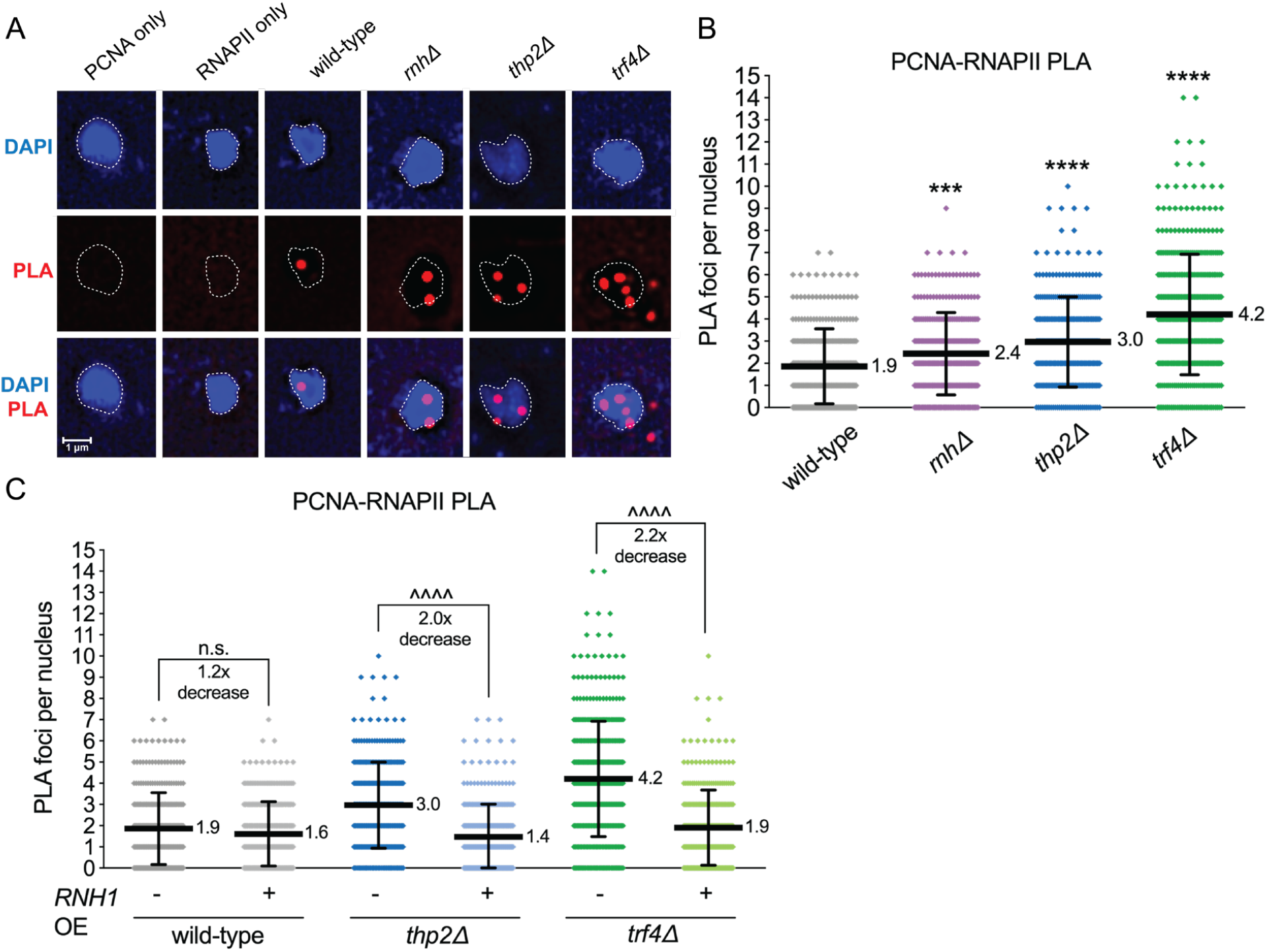
Transcription-replication conflicts (TRCs) in RNA biogenesis mutants. (A) Example images for the Proximity Ligation Assay (PLA). DAPI staining the nucleus is shown in blue, PLA foci (proximity of PCNA and RNAPII-pSer2) shown in red. The first two columns are single-antibody control conditions and the other columns with both antibodies. (B) PLA using an antibody to RNAPII-pSer2 and one to PCNA to assess TRCs genome-wide in the indicated strains. N≥300 nuclei quantified per condition. Dots indicated individual PLA foci counts per nucleus. Error bars show mean ± SD. ***p<0.001 or ****p<0.0001 compared to wild-type by Mann-Whitney test (Table S9). (C) PLA assessing TRCs upon RNase H1 overexpression. N≥300 nuclei quantified per condition. Dots indicated individual PLA foci counts per nucleus. Error bars show mean ± SD. ^^^^p<0.0001 compared to indicated strain by Mann-Whitney test (Table S9).

To further understand the basis of the TRCs, we analyzed numbers of PLA foci in the strains with endogenously inducible RNase H1. Overexpression of RNase H1 reduced PLA foci numbers in both *thp2Δ* and *trf4Δ* to wild-type levels, (Fig. 4C, Table S9). There was no significant reduction in PLA foci upon RNase H1 overexpression in a wild-type strain (Fig. 4C, Table S9). These data suggest that RNase H1 resolves RNA:DNA hybrids that form at the sites of conflicts between RNAPII and a replication fork in THO and TRAMP4 mutants.

### Cells lacking Trf4 exhibit an RPA binding deficiency that leads to CAG repeat fragility and TRCs

Since RNase H1 overexpression did not rescue CAG fragility in the *trf4Δ* mutant, we investigated other possible reasons for the increased breaks that occur in the absence of TRAMP4. RPA recruitment is decreased in the absence of Trf4 or Rrp6 after resection of DSBs, and this attenuates the Mec1/ATR response to DSBs as well as to fork-stalling agents HU and MMS [61]. *Trf4Δ* does not affect resection of an HO-induced break [61]. Additionally, the catalytic RNA degradation activity of the mammalian Rrp6 homolog EXOSC10 is needed for normal levels of RPA recruitment to sites of irradiation-induced DSBs [62]. RPA-loading is expected to be crucial for preventing hairpin formation at single-stranded CAG or CTG repeats [63], and indeed RPA was shown to accumulate at CAG repeats by induction of convergent transcription [64]. To test if RPA recruitment was impaired when RNA degradation factors were deleted, we performed ChIP to detect levels of RPA at the CAG repeat using an RPA antibody that recognizes all three yeast RPA subunits. We observed a 1.2-fold increase in enrichment of RPA proteins at the CAG-70 repeat over a locus within the *ACT1* gene (*ACT1* locus) (Fig. 5A), indicating that more RPA is recruited to the expanded CAG repeat tract than a regular genomic locus. The relative RPA ChIP signal at the CAG-70 repeats in the *trf4Δ* strain was reduced significantly by ∼30% compared to the wild-type control (p= 0.03) (Fig. 5A, Table S8). The RPA ChIP level in the *trf4Δthp2Δ* double mutant showed a similar 30% level of reduction compared to wild-type (p=0.05), strengthening this conclusion. In contrast, RPA is still enriched at the CAG repeat tract in the *thp2Δ* mutant (Fig. 5A). These data indicate that when lacking Trf4, RPA association at the CAG-70 repeat is diminished.

**Figure 5.**
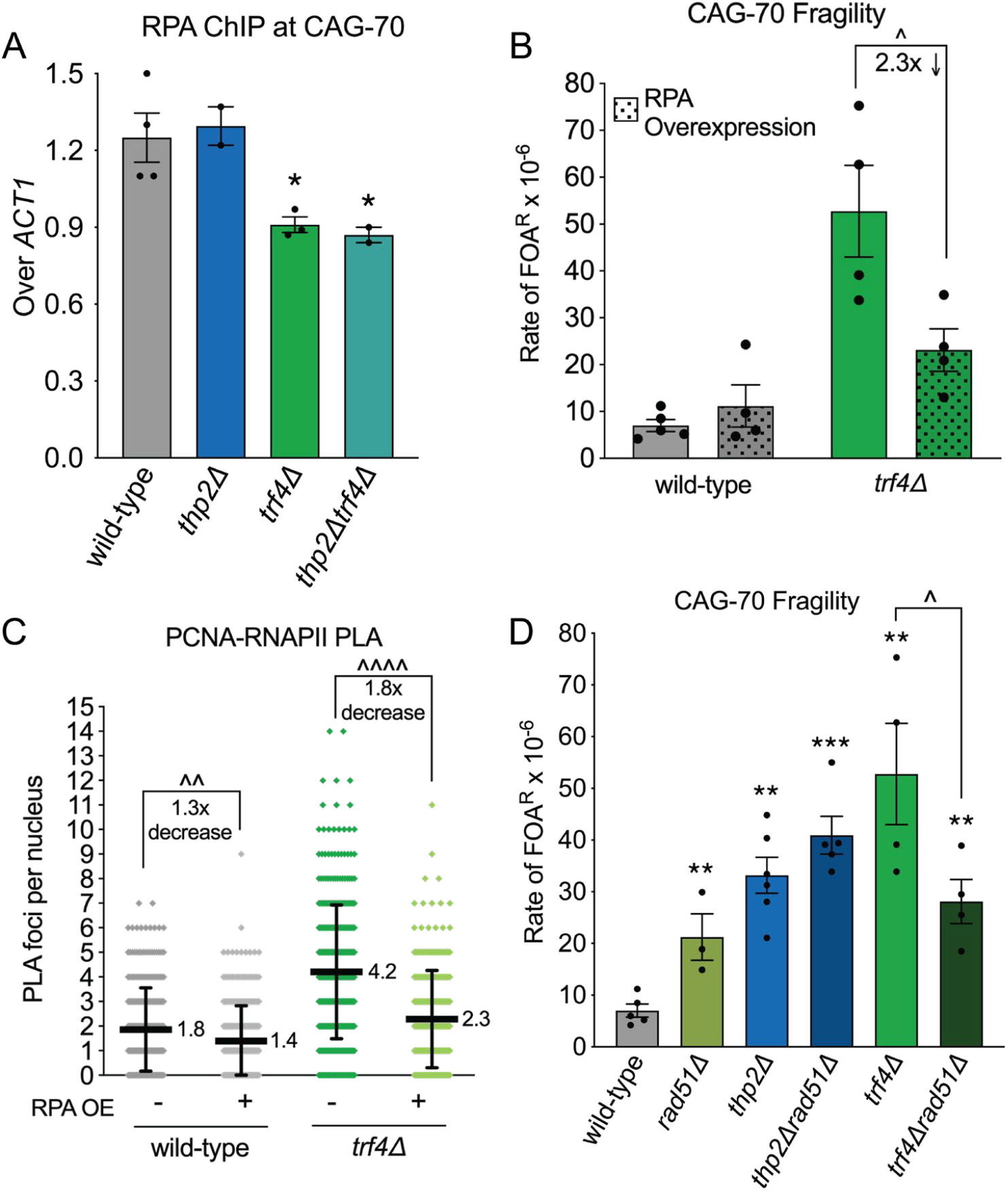
RPA-loading deficiency and genetic interactions in the *trf4Δ* strain. (A) RPA chromatin immunoprecipitation (ChIP) at CAG repeats in the indicated strains. Each bar represents the mean ± SEM of at least two experiments from two biological replicates and each dot represents an individual biological replicate; *p<0.05, compared to wild-type by *t*-test (Table S8). (B) Rate of FOA^R^ × 10^−6^ in indicated mutants and RPA overexpression conditions (strains transformed with RPA overexpression vector compared to no vector). Each dot represents an individual data point; ^p<0.05, compared to no induction condition in the same mutant, by *t*-test. Average of at least 3 experiments ± SEM is shown (Table S1). (C) PLA assessing TRCs in strains with and without RPA overexpression. N≥300 nuclei quantified per condition. Dots indicated individual PLA foci counts per nucleus. Error bars show mean ± SD. ^^^^p<0.0001 compared to indicated strain by Mann-Whitney test (Table S9). (D) Rate of FOA^R^ × 10^−6^ in indicated mutants; each dot represents an individual fragility experiment; **p<0.01 and ***p <0.001, compared to wild-type with same CAG tract, ^p=0.06, compared to *trf4Δ* by *t*-test. Average of at least 3 experiments ± SEM is shown (Table S1).

A reduced level of RPA binding is predicted to increase the chance of hairpin formation in exposed ssDNA regions, which could interfere with DNA replication or cause inefficient repair, increasing the chance of breakage or failure to repair, resulting in YAC end loss. To test whether the reduced RPA binding observed at the CAG tract in the *trf4Δ* strain was contributing to the increase in fragility, we overexpressed RPA by introducing a multicopy plasmid containing all three RFA genes [65]. This led to a 20-40-fold increase in *RFA1, RFA2*, and *RFA3* expression over basal levels (Fig S2B, Table S11). We observed a significant suppression of CAG tract fragility upon overexpression of RPA in the *trf4Δ* strain (44% reduction, 2.3-fold decrease) that was not observed in wild-type cells (Fig. 5B, Table S1). This result indicates that the reduced RPA binding observed at the CAG tract in the *trf4Δ* strain may be the source of the increased repeat tract fragility in the absence of TRAMP4 activity.

To determine whether RPA levels were having a genome-wide consequence when the TRAMP4 complex is defective we tested whether RPA overexpression could relieve the increased level of TRCs observed. Indeed, TRCs in the *trf4Δ* strain were suppressed to wild-type levels upon RPA overexpression (1.8-fold decrease, p<0.0001; Fig. 5C, Table S9). This result indicates that lack of RPA availability leads to the increased TRCs observed in the absence of TRAMP4. Though less dramatic, TRCs were also reduced upon RPA overexpression in wild-type cells (1.3-fold, p=0.001; Fig. 5C, Table S9), indicating that RPA may also be limiting at spontaneous TRCs that occur, impacting their resolution.

Finally, to determine whether strand annealing activity was required to recover from the TRCs observed in RNA biogenesis mutants and prevent fragility, we tested the role of the Rad51 protein. CAG-70 fragility levels were similar for *thp2Δ* and *thp2Δ*r*ad51Δ* cells, indicating that the damage occurring at the CAG tract in THO mutants is generally not rescued by homologous recombination (Fig 5D, Table S1). Surprisingly, fragility was reduced in the *trf4Δ*r*ad51Δ* mutant (Fig 5D, Table S1). Therefore, the presence of Rad51 is exacerbating CAG fragility in cells lacking TRAMP4. In situations where forks are de-protected or cannot restart, excess Rad51 binding can lead to deleterious accumulation of recombination structures at stalled forks [66]. Therefore, the suppressive effect of deleting Rad51 in the *trf4Δ* strain may be due to RPA depletion at CAG stalled forks that allows for excessive Rad51 binding and formation of deleterious recombination intermediates, leading to increased fragility.

## Discussion

In this study, we discovered that CAG repeat fragility is greatly increased in the absence of components of the THO or TRAMP complexes or the exosome, indicating that defects in RNA biogenesis and processing lead to frequent occurrence of chromosome breakage events. We examined several potential causes of the fragility in the Thp2 (THO) and Trf4 (TRAMP4) mutants, including R-loop accumulation, RNA polymerase stalling, TRCs, and defects in RPA recruitment. We found that RNAPII stalling at expanded CAG repeats was the most prominent phenotype for both *thp2Δ* and *trf4Δ* mutants and is likely the main initiator of the increased CAG fragility. Supporting this conclusion, genome-wide TRCs were significantly increased in both *thp2Δ* and *trf4Δ* mutants, to a much greater degree than a strain missing both RNase H1 and RNase H2 that has a confirmed increase in RNA:DNA hybrids. Consistent with our *in vivo* data, a recent *in vitro* study using *E. coli* proteins showed that RNA polymerase transcription complexes, especially if oriented head-on with replication, created stable blockages that were more severe than an R-loop without an attached RNA polymerase [67]. Based on the 7.0-fold greater level of transcription in the rCUG orientation at the CAG repeat studied here [7], we predict that most conflicts within the CAG tract will be in the head-on orientation. Altogether, our results show that defects in both RNP formation on nascent RNA and RNA processing lead to RNAPII stalling and TRCs, and that chromosome fragility is a frequent and deleterious consequence.

Since the THO-defective *hpr1Δ* mutant was shown to exhibit increased R-loop formation and TAR [22], and R-loop accumulation in RNase H mutants increases CAG repeat fragility [7], we predicted that R-loops would be the main cause of fragility at the expanded CAG tract in the THO mutant. Unexpectedly, we were not able to detect increased R-loop signals at the CAG tract in the *thp2Δ* mutant by DRIP as we did for RNase H depletion, though we did see a partial but significant decrease in CAG fragility when RNase H1 was overexpressed in the *thp2Δ* strain. Also, TRCs were dramatically reduced by *in vivo* RNase H1 overexpression. Because we could not directly detect an increase in R-loops at the CAG tract by DRIP in the *thp2Δ* mutant, but only detected their phenotypic consequence by the RNase H1 overexpression experiments, we hypothesize that they are transient and likely associated with the stalled RNAPII, rather than stable R-loops left after passage of the transcriptional machinery. It has been recently recognized that small R-loops (60 bp or less) associated with paused RNAPII are not detected by DRIP, which preferentially detects longer (>200 bp), more stable R-loops [55]. Based on the role of THO in binding nascent RNA to direct it to the nuclear pore complex for export, it is probable that the unbound nascent RNA is more likely to re-anneal to the DNA template in this mutant, inhibiting RNAPII progression and causing an accumulation of RNAPII within the CAG tract (Fig. 6, left pathway). Indeed, THO is the complex that binds closest to RNAPII and has a direct role in transcription elongation [19]. Since overexpression of RNase H1 decreased TRCs in the *thp2Δ* mutant, these tethered RNA polymerases likely block the replication fork, leading to TRCs, fragility, and repeat contractions (Fig. 6). Digestion of the RNA of a co-transcriptional RNA:DNA hybrid could release stalled RNAPII, leading to the observed suppression of TRCs and breaks.

**Figure 6.**
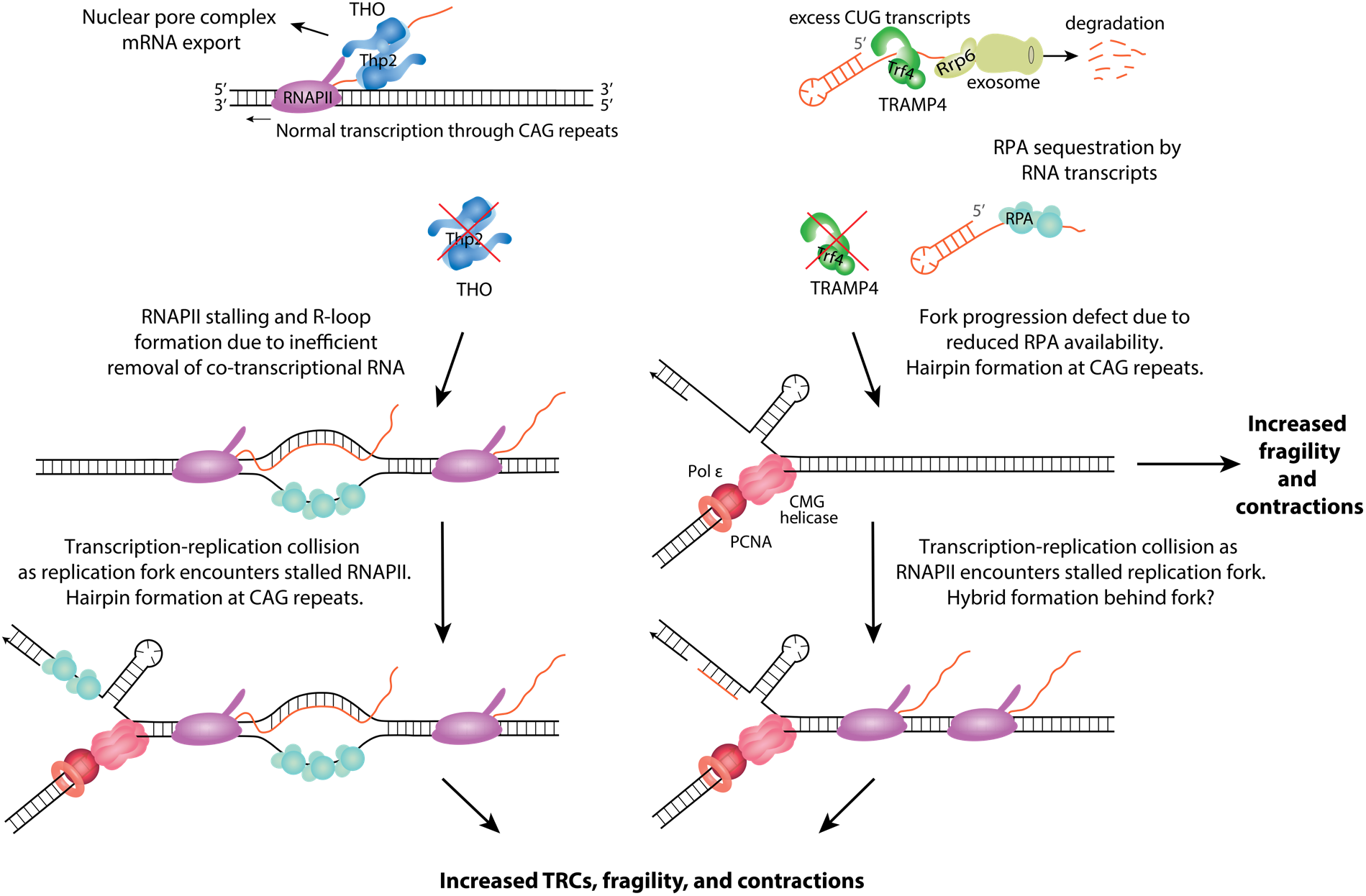
Model for CAG repeat fragility arising from defective THO and TRAMP4 complexes. Transcription through CAG repeats is shown (Top), in the direction of the read-through transcription from the *URA3* gene on the YAC (rCUG transcript), though some cryptic transcription occurs from the other direction as well (rCAG transcript). Under wild-type conditions, RNAPII generates CAG or CUG transcripts, which are co-transcriptionally bound by THO (blue). They may be recognized as cryptic unstable transcripts (CUTs), bound and polyadenylated by TRAMP4 (green), and targeted for exosome-mediated degradation through Rrp6 and the exosome (olive green). In the absence of THO (left), RNAPII stalling and R-loop formation occurs due to inefficient removal of nascent RNA that hybridizes to the exposed ssDNA. In this scenario, RNAPII stalling is the initiating event. The increased RNAPII stalling leads to transcription-replication conflicts (TRCs) as the replication fork approaches and encounters stalled RNAPII. The stalled replication fork and exposed ssDNA allows for CAG hairpin formation, leading to fragility and contractions of the CAG repeats. In the absence of TRAMP4 (right), a replication fork progression defect occurs due to impaired RPA loading, allowing for hairpin formation at the CAG repeats and replication fork stalling, leading to fragility and contractions. In this scenario, replication fork stalling is the initiating event. TRCs occur as RNAPII encounters the stalled replication fork, potentially accompanied by short RNA:DNA hybrid formation. This ultimately leads to additional fragility and contractions of the CAG repeats.

Our results with the Thp2 mutant highlight the importance of successful transcription elongation in preventing TRCs and chromosome breaks not only within genes, but also in repetitive DNA. Mutation of other proteins involved in mRNA maturation also have genome protective effects. A mutation in Ysh1, an endoribonuclease subunit of the mRNA cleavage and polyadenylation complex, resulted in slowed RNAPII passage through a GAA repeat tract, and GAA expansions and fragility in an R-loop-independent manner [68]. Excess Yra1, a member of the mRNA export TREX complex, causes genome-wide replication slowing and increased DNA damage, ultimately causing telomere shortening and replicative senescence [69, 70]. Thus, interference at multiple points of the RNA maturation process can lead to similar outcomes. A recent study showed that human THOC7, a component of human THO, accumulates in transcriptionally active repeat regions and defects are associated with γH2AX foci, suggesting that our results will be relevant to maintenance of repeats in the human genome [71].

The TRAMP complexes target mainly non-coding RNAs to Rrp6 and the nuclear exosome for degradation, a pathway that is highly conserved from yeast to humans. It was previously shown that *trf4Δ* mutants exhibit increased transcription-dependent recombination and an elevated mutation rate that was linked to the presence of excess nascent RNA transcripts [36]. These genome instability phenotypes were attributed to increased R-loops since they were reduced upon overexpression of RNase H1 [36]. Strains depleted of Trf4 or Rrp6 showed increased RNase H1-dependent chromosome loss and terminal deletions [37], which were suggested to be caused by in-trans R-loops that are dependent on Rad51 and Rad52 [40]. Rad51-dependent accumulation of TERRA RNA at telomeres is also increased in *trf4Δ* strains [72]. However, at the CAG tract, we could find no evidence of increased R-loops either by DRIP or RNase H1 overexpression experiments. Instead, we found a significant increase in RNAPII accumulation over the CAG tract and a high level of genome-wide TRCs in the *trf4Δ* mutant. Therefore, we explored whether the primary cause of CAG fragility and TRCs in the absence of a functional TRAMP4 complex could be related to RPA levels on DNA. Based on our observation of a decrease in RPA binding to CAG repeats and a suppression of both CAG repeat fragility and TRCs upon RPA overexpression in the *trf4Δ* mutants, we propose an alternative hypothesis for genome instability phenotypes in the absence of TRAMP4 (Fig. 6, right). A deficiency of RPA binding to replication forks paused within the CAG tract could allow more hairpin formation or could expose naturally stalled forks to excess degradation, either of which could result in decreased fork recovery. This model is consistent with the observed suppression in CAG repeat fragility and genome-wide TRCs upon overexpression of RPA in the *trf4Δ* mutant. It can also explain the suppression of fragility by loss of Rad51, since a fork unprotected by RPA binding will be more available for Rad51 loading and unregulated HR, which could lead to cleavage or failure to effectively restart. At sites of R-loop-mediated TRCs, RECQ5 is needed to disrupt RAD51 filaments to allow for fork restart that is mediated by MUS81 cleavage [73].

The question remains of why RPA levels on DNA are reduced in TRAMP4 mutants. TRAMP mutants have an excess accumulation of cryptic RNA [32, 34, 36, 42]. A recent study showed that RPA can bind with high affinity to ssRNA [74]. Therefore, RPA loading at stalled forks could be decreased due to excess unprocessed RNAs in the nucleoplasm of *trf4Δ* cells competing for RPA binding. Another consideration is that RNA:DNA hybrids appear to contribute to the TRCs observed in the *trf4Δ* mutant as we observed a suppression of TRCs upon overexpression of RNase H1, indicating that degradation of hybrids could allow for TRC resolution. It has been recently proposed that hybrids form at stalled forks and interfere with fork restart [75, 76]. It is possible that low levels of RPA due to sequestration by excess RNAs in the absence of TRAMP4 favors formation of hybrids behind the fork, inhibiting TRC resolution (Fig. 6, right). Suggesting conservation, in EXOSC10 (yeast Rrp6) depleted human cells, RPA recruitment to a DSB was restored upon clearance of damage-induced long non-coding RNAs (dilncRNAs) by treatment with RNase H1 [62]. Altogether, our data support that the TRAMP4 complex plays an important role in preventing TRCs and chromosomal breaks through processing of cryptic RNAs and preventing their accumulation and sequestration of RPA.

Even though the THO and TRAMP complexes are both involved in RNA biogenesis, we found that deletion of both pathways was highly synergistic for CAG fragility. In addition, deletion of Trf4 along with RNase H1 and RNase H2 was strongly synergistic, consistent with our conclusion that *trf4* mutants cause CAG fragility by a pathway not limited to increasing R-loops. The *thp2ΔrnhΔ* double mutant also showed synergistic fragility, though less than the other combinations, in line with THO mutants causing CAG fragility through both R-loop and non-R-loop mechanisms. Our results indicate that RNA biogenesis mutants can cause genome instability by multiple mechanisms that are only partially overlapping, including interference with RNAPII progression or unloading, TRCs, R-loops, and reduced RPA binding to DNA. Since the fragility phenotypes were not just additive but synergistic, it implies that the activities of these protein complexes work in a cooperative manner to prevent genome instability. The conversion of TRCs to a double-strand break could occur by several mechanisms and the downstream events that lead to chromosome fragility will be an interesting area of future investigation.

In summary, our results highlight the importance of multiple aspects of RNA biogenesis in preventing genome stability and show that these mechanisms are especially crucial at structure-forming repeats. Since repetitive DNA occurs throughout genomes, these pathways are expected to be of paramount importance in preventing breaks and the ensuing deleterious consequences.

## Materials and Methods

### Yeast Strains and Genetic Manipulation

Yeast strains used in this study are listed in Table S13. Yeast knock-out mutants were created by one-step gene replacement [77] using selectable markers, *KANMX6, TRP1*, or *HIS3MX6* and method described in [77]. The Met25 promoter was inserted right upstream of the *RNH1* gene in different strain backgrounds by homology-directed replacement; the construct of the *MET25* promoter with a *natNT2* marker gene is from pYM-N35 [53]. Overexpression of RPA was achieved by transforming desired strains with a multicopy plasmid containing *RFA1, RFA2*, and *RFA3* [65]. PCR was used to confirm the successful knockout of a gene by confirming the presence of the selectable marker and the absence of the endogenous gene at the target locus. CAG tract length was verified in all successful clones by PCR as described below.

### CAG Repeat Tract Length Verification and Fragility Analysis

The CAG tract was verified by using PCR amplification of genomic DNA from yeast colonies using colony PCR with primers specific to the CAG repeat tract listed in Table S12. The PCR protocol was described in [44] and the product sizes were analyzed by high-resolution gel electrophoresis. 10 colonies carrying the correct length CAG tract were used in one fragility assay as described in [44]. See also [78, 79] for a detailed protocol. The colonies grown on FOA-Leu and YC-Leu were counted, and the rate of FOA resistance was calculated by using the Ma-Sandri-Sarkar Maximum Likelihood Estimator (MSS-MLE) [80, 81]. The loss of the *URA3* marker on the YAC for at least 10 independent FOA-resistant colonies was confirmed by either Southern blot or PCR for the wild-type, *thp2Δ*, and *trf4Δ* mutants. Due to high FOA-resistance and repeat contraction frequencies in the *thp2Δtrf4Δ* mutant, a regular 10-colony fragility assay and the MSS-Maximum Likelihood estimation were not applicable. Therefore, a one-colony fragility assay was carried out. Each assay was from one parental colony with desired CAG-70 tract length. The mutant cells were grown in YC-Leu media for an extended time (∼48 hours) for ∼5 doublings instead of 6-7 divisions in regular assays. Then, the FOA-resistant frequency was calculated by counting the cells grown on the FOA-Leu and YC-Leu plates.

### DNA:RNA Immunoprecipitation (DRIP) and Chromatin Immunoprecipitation (ChIP)

DRIP was performed by using the same procedure as described in [7]. S9.6 (4μg, Kerafast) antibody [82, 83] was used to coat 40 μL of Protein G Dynabeads (Invitrogen) per sample. At least two, but usually three biological replicates were done for each condition. RPA and RNAPII ChIP were done using the same procedure as the ChIP described in [7], except in the antibodies used. Unsynchronized cells were cross-linked in 1% formaldehyde for 20 minutes at room temperature. Antibody usage is as follows; anti-RPA (Agisera): 5 uL of 1:4 dilution from original stock for each sample (concentration undetermined by the company due to its serum format); anti-RNA PolII pSer5, raised against 10 repeats of YSPTSPS peptide (CTD4H8, Santa Cruz): 1 μg for each sample. IP and input DNA levels were quantified by qPCR using SYBR green PCR mastermix (Roche) on a 7300 real-time PCR system (Applied Biosystems) or using SYBR Premix Ex Taq II (Tli RNase H Plus) (Takara Bio) on a LightCycler 480 II (Roche). qPCR reactions were performed in technical duplicate.

### Quantitative reverse transcription PCR (qRT-PCR)

The RNA preparation and reverse transcription procedure used is that described in [12], with the Illustra RNAspin mini kit (GE Healthcare) or RNeasy kit and RNase-free DNase set (QIAGEN) and Superscript First Strand Synthesis kit (Life Technologies); oligo d(T) primers were used for priming during reverse transcription. RT-PCR samples were analyzed using qPCR with SYBR green PCR mastermix (Roche), SYBR Premix Ex Taq II (Tli RNase H Plus) (Takara Bio), or Power SYBR Green PCR Master Mix (Applied Biosystems) on 7300 real-time PCR system (Applied Biosystems), LightCycler 480 II (Roche), or QuantStudio 6 real-time PCR system (Applied Biosystems).

### Proximity Ligation Assay

Proximity ligation assays (PLA) were performed using the Duolink kit from Millipore Sigma. Preparation of yeast cells was adapted from [84]. Cells were grown in YC-Ura-Leu media (YC-Ura-Leu-Met-Cys media for RNase H overexpression and YC-His-Ura-Leu for RPA overexpression PLA experiments) at 30°C to log phase. Cells were fixed with 4% paraformaldehyde for 15 min at room temperature. Cells were washed 3x with wash buffer (1.2M sorbitol in 100 mM KPO_4_ pH 6.5). Cell walls were digested in zymolyase solution (500 μg/mL Zymolyase 100T, 1.2M sorbitol, 100 mM KPO_4_ pH 6.5, 20 mM 2-Mercaptoethanol) at 30°C shaking for 30 min. Cells were washed 3x with wash buffer and resuspended in 1.2M sorbitol. Cells were attached to poly-L-lysine coated slides and washed 3x with wash buffer and 3x with permeabilizing solution (1% TritonX-100 in 100 mM KPO_4_ pH 6.5). Cells were blocked with the provided blocking solution for 30 min at 37°C (Duolink kit). Cells were incubated with primary antibody (1:400 RNAPII [pSer2] Novus Biologicals NB100-1805, 1:400 PCNA [5E6/2] abcam ab70472) overnight at 4°C. Cells were washed 2× in wash buffer A (Duolink kit) and incubated in anti-rabbit PLUS and anti-mouse MINUS probes diluted 1:5 in antibody diluent (Duolink kit) for 1 hr at 37°C. Cells were washed 2× in wash buffer A and incubated in ligase solution (1:40 ligase in 5× ligase buffer diluted 1:5 in water, Duolink kit) for 30 min at 37°C. Cells were washed 2× in wash buffer A and incubated in amplification solution (1:80 polymerase in 5× amplification buffer diluted 1:5 in water, Duolink kit) for 100 min at 37°C. Cells were washed 2× in wash buffer B (Duolink kit) and 1× in 0.01× wash buffer B. The coverslip was mounted with In Situ Mounting Medium with DAPI (Duolink kit) and sealed with nail polish. Slides were imaged using a Leica Dmi8 Thunder or a DeltaVision Ultra microscope at 100× oiled objective. Number of PLA foci per DAPI-stained nucleus was quantified.

## Supplementary Figures

**Figure S1.**
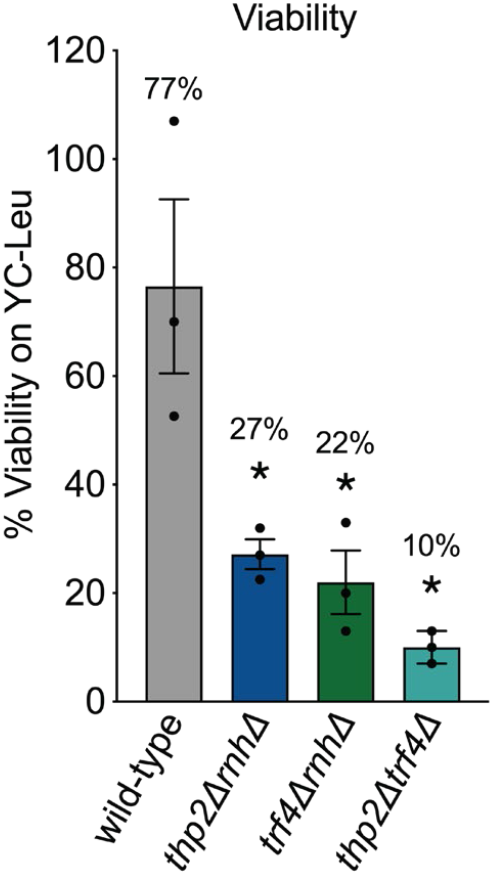
Cell viability in mutant backgrounds. Frequencies of viability are shown. Viability is calculated by comparing the amount of cells that grew on YC-Leu to the amount of cells counted by hemacytometer. Each dot represents an individual data point; average ± SEM is shown; *p<0.05, compared to wild-type, *t* test (Table S5).

**Figure S2.**
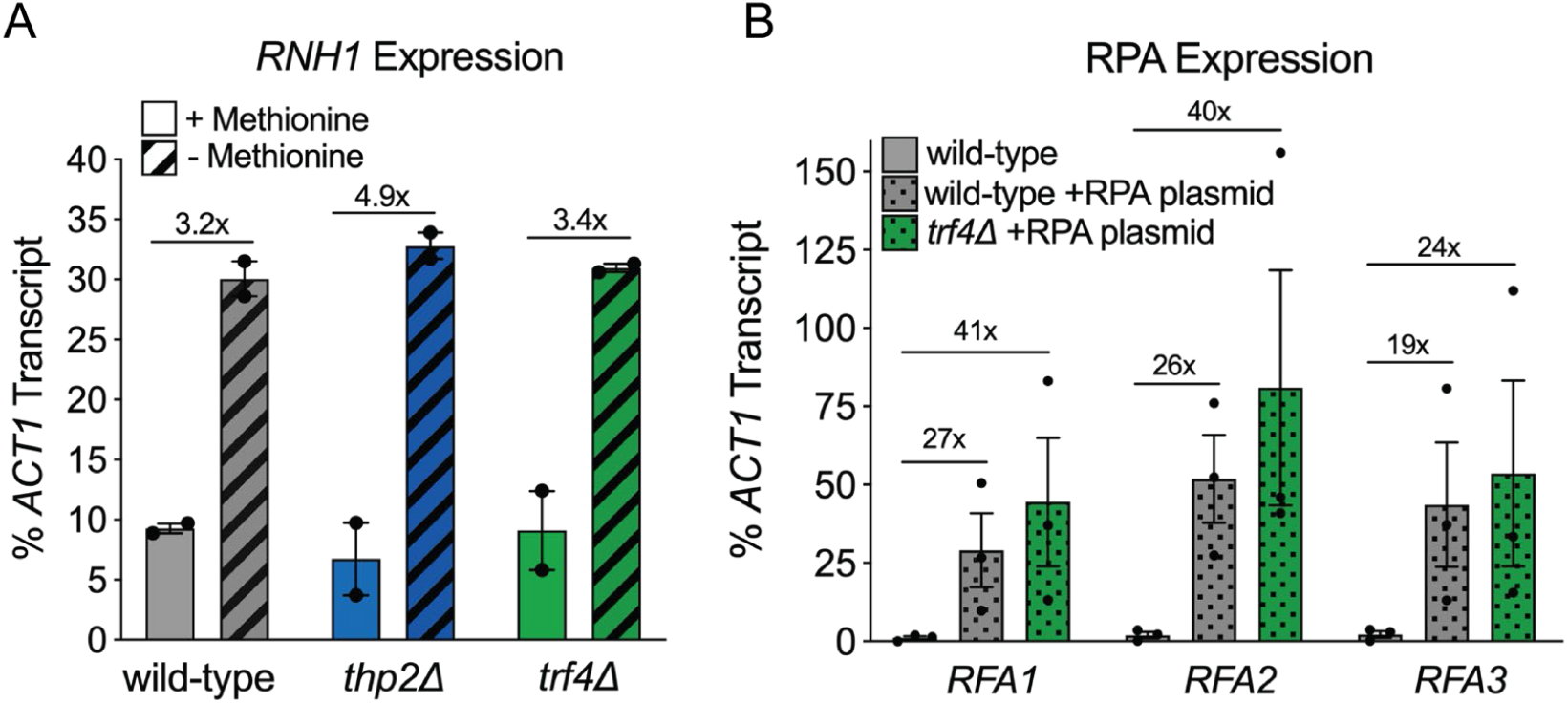
*RNH1* and RPA transcript levels. (A) Overexpression of *RNH1* gene under Met25 promoter is induced in the media lacking methionine and cysteine. mRNA was reversed transcribed into cDNA by RT-PCR and qPCR was used to quantify cDNA at *RNH1* and *ACT1* gene loci. *RNH1* (under Met25 promoter) qPCR signal was normalized to the *ACT1* (under endogenous promoter) qPCR signal in the indicated mutants (Table S7). (B) Overexpression of RPA by transforming strains with multicopy plasmid containing *RFA1, RFA2*, and *RFA3*. mRNA was reversed transcribed into cDNA by RT-PCR and qPCR was used to quantify cDNA at *RFA1/RFA2/RFA3* and *ACT1* gene loci. *RFA1/RFA2/RFA3* were normalized to the *ACT1* (under endogenous promoter) qPCR signal in the indicated strains (Table S11).

**Figure S3:**
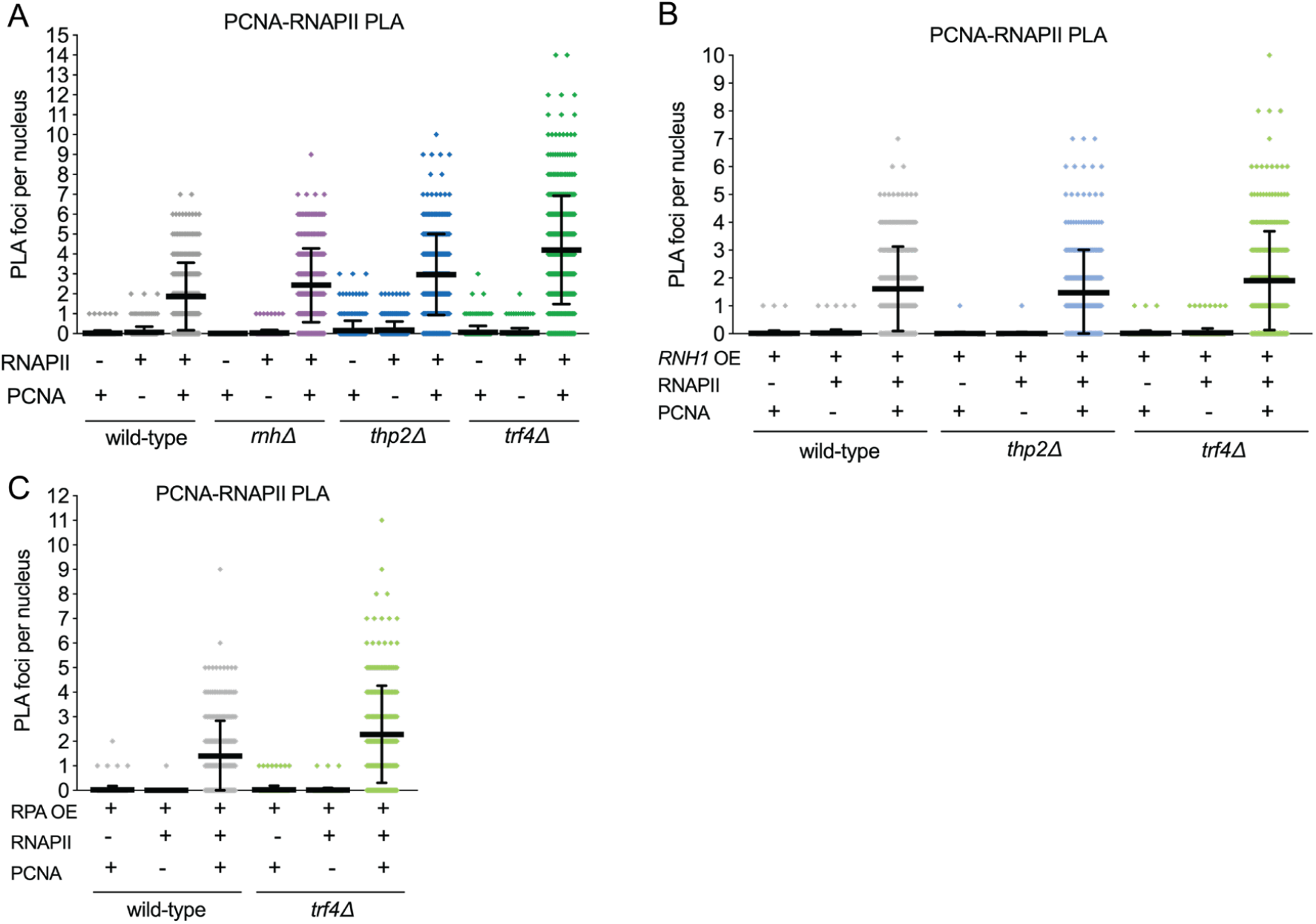
Proximity Ligation Assay (PLA) single antibody controls. Antibodies to the Ser2 phosphorylated form of RNAPII (RNAPII-pSer2) and one to PCNA were used to assess transcription-replication conflicts (TRCs) genome-wide in the indicated strains. N≥300 nuclei quantified per condition. Dots indicated individual PLA foci counts per nucleus. Error bars show mean ± SD. See Table S9 for p-values of single antibody controls compared to double antibody conditions, by Mann-Whitney test. Double antibody (RNAPII and PCNA) experiments are shown alongside single antibody controls (Table S9). (A) PLA in wild-type, *rnh1Δrnh201Δ, thp2Δ*, and *trf4Δ* strains. (B) PLA in wild-type, *thp2Δ*, and *trf4Δ* strains with RNase H1 overexpression. (C) PLA in wild-type and *trf4Δ* strains with RPA overexpression.

## Acknowledgements

We thank Cailin Joyce, who re-tested *THP2* in the original CAG fragility screen, as well as the students in the 2015 and 2017 Tufts Molecular Genetics Project Lab who identified *TRF4, MFT1*, and *LRP1* genes. We also thank Meaghan McGoldrick who helped with strain construction and Alexandra Khristich and Sergei Mirkin for sharing the RPA overexpression plasmid.

## Funding

This work was funded by NSF MCB 1330743 and NSF MCB 1817499 awarded to C.H.F.

## Notes

### Competing Interest Statement

The authors have declared no competing interest.

## References

1. Usdin K, House NC, Freudenreich CH. Repeat instability during DNA repair: Insights from model systems. Crit Rev Biochem Mol Biol. 2015;50(2):142–67. Epub 2015/01/23. doi: 10.3109/10409238.2014.999192. PubMed PMID: 25608779; PubMed Central PMCID: PMCPMC4454471.

2. Brown RE, Freudenreich CH. Structure-forming repeats and their impact on genome stability. Curr Opin Genet Dev. 2020;67:41–51. Epub 2020/12/07. doi: 10.1016/j.gde.2020.10.006. PubMed PMID: 33279816.

3. Khristich AN, Mirkin SM. On the wrong DNA track: Molecular mechanisms of repeat-mediated genome instability. J Biol Chem. 2020;295(13):4134–70. Epub 2020/02/16. doi: 10.1074/jbc.REV119.007678. PubMed PMID: 32060097; PubMed Central PMCID: PMCPMC7105313.

4. Goula AV, Stys A, Chan JP, Trottier Y, Festenstein R, Merienne K. Transcription elongation and tissue-specific somatic CAG instability. PLoS Genet. 2012;8(11):e1003051. doi: 10.1371/journal.pgen.1003051. PubMed PMID: 23209427; PubMed Central PMCID: PMCPMC3510035.

5. Goula AV, Pearson CE, Della Maria J, Trottier Y, Tomkinson AE, Wilson DM, 3rd, et al. The nucleotide sequence, DNA damage location, and protein stoichiometry influence the base excision repair outcome at CAG/CTG repeats. Biochemistry. 2012;51(18):3919–32. doi: 10.1021/bi300410d. PubMed PMID: 22497302; PubMed Central PMCID: PMCPMC3357312.

6. Lin Y, Wilson JH. Transcription-induced CAG repeat contraction in human cells is mediated in part by transcription-coupled nucleotide excision repair. Mol Cell Biol. 2007;27(17):6209–17. doi: 10.1128/MCB.00739-07. PubMed PMID: 17591697; PubMed Central PMCID: PMCPMC1952160.

7. Su XA, Freudenreich CH. Cytosine deamination and base excision repair cause R-loop-induced CAG repeat fragility and instability in Saccharomyces cerevisiae. Proc Natl Acad Sci U S A. 2017;114(40):E8392–E401. doi: 10.1073/pnas.1711283114. PubMed PMID: 28923949; PubMed Central PMCID: PMCPMC5635916.

8. Lin Y, Dent SY, Wilson JH, Wells RD, Napierala M. R loops stimulate genetic instability of CTG.CAG repeats. Proc Natl Acad Sci U S A. 2010;107(2):692–7. doi: 10.1073/pnas.0909740107. PubMed PMID: 20080737; PubMed Central PMCID: PMCPMC2818888.

9. Reddy K, Schmidt MH, Geist JM, Thakkar NP, Panigrahi GB, Wang YH, et al. Processing of double-R-loops in (CAG).(CTG) and C9orf72 (GGGGCC).(GGCCCC) repeats causes instability. Nucleic Acids Res. 2014;42(16):10473–87. doi: 10.1093/nar/gku658. PubMed PMID: 25147206; PubMed Central PMCID: PMCPMC4176329.

10. Reddy K, Tam M, Bowater RP, Barber M, Tomlinson M, Nichol Edamura K, et al. Determinants of R-loop formation at convergent bidirectionally transcribed trinucleotide repeats. Nucleic Acids Res. 2011;39(5):1749–62. doi: 10.1093/nar/gkq935. PubMed PMID: 21051337; PubMed Central PMCID: PMCPMC3061079.

11. Laverde EE, Lai Y, Leng F, Balakrishnan L, Freudenreich CH, Liu Y. R-loops promote trinucleotide repeat deletion through DNA base excision repair enzymatic activities. J Biol Chem. 2020;295(40):13902–13. Epub 2020/08/09. doi: 10.1074/jbc.RA120.014161. PubMed PMID: 32763971; PubMed Central PMCID: PMCPMC7535915.

12. Koch MR, House NCM, Cosetta CM, Jong RM, Salomon CG, Joyce CE, et al. The Chromatin Remodeler Isw1 Prevents CAG Repeat Expansions During Transcription in Saccharomyces cerevisiae. Genetics. 2018;208(3):963–76. doi: 10.1534/genetics.117.300529. PubMed PMID: 29305386.

13. Xu J, Chong J, Wang D. Strand-specific effect of Rad26 and TFIIS in rescuing transcriptional arrest by CAG trinucleotide repeat slip-outs. Nucleic Acids Res. 2021;49(13):7618–27. Epub 2021/07/02. doi: 10.1093/nar/gkab573. PubMed PMID: 34197619; PubMed Central PMCID: PMCPMC8287942.

14. Salinas-Rios V, Belotserkovskii BP, Hanawalt PC. DNA slip-outs cause RNA polymerase II arrest in vitro: potential implications for genetic instability. Nucleic Acids Res. 2011;39(17):7444–54. Epub 2011/06/15. doi: 10.1093/nar/gkr429. PubMed PMID: 21666257; PubMed Central PMCID: PMCPMC3177194.

15. Burns JA, Chowdhury MA, Cartularo L, Berens C, Scicchitano DA. Genetic instability associated with loop or stem-loop structures within transcription units can be independent of nucleotide excision repair. Nucleic Acids Res. 2018;46(7):3498–516. Epub 2018/02/24. doi: 10.1093/nar/gky110. PubMed PMID: 29474673; PubMed Central PMCID: PMCPMC5909459.

16. Luna R, Rondon AG, Aguilera A. New clues to understand the role of THO and other functionally related factors in mRNP biogenesis. Biochim Biophys Acta. 2012;1819(6):514–20. Epub 2011/12/31. doi: 10.1016/j.bbagrm.2011.11.012. PubMed PMID: 22207203.

17. Chavez S, Beilharz T, Rondon AG, Erdjument-Bromage H, Tempst P, Svejstrup JQ, et al. A protein complex containing Tho2, Hpr1, Mft1 and a novel protein, Thp2, connects transcription elongation with mitotic recombination in Saccharomyces cerevisiae. EMBO J. 2000;19(21):5824–34. doi: 10.1093/emboj/19.21.5824. PubMed PMID: 11060033; PubMed Central PMCID: PMCPMC305808.

18. Pena A, Gewartowski K, Mroczek S, Cuellar J, Szykowska A, Prokop A, et al. Architecture and nucleic acids recognition mechanism of the THO complex, an mRNP assembly factor. EMBO J. 2012;31(6):1605–16. Epub 2012/02/09. doi: 10.1038/emboj.2012.10. PubMed PMID: 22314234; PubMed Central PMCID: PMCPMC3321177.

19. Gewartowski K, Cuellar J, Dziembowski A, Valpuesta JM. The yeast THO complex forms a 5-subunit assembly that directly interacts with active chromatin. Bioarchitecture. 2012;2(4):134–7. Epub 2012/09/12. doi: 10.4161/bioa.21181. PubMed PMID: 22964977; PubMed Central PMCID: PMCPMC3675074.

20. Zenklusen D, Vinciguerra P, Wyss JC, Stutz F. Stable mRNP formation and export require cotranscriptional recruitment of the mRNA export factors Yra1p and Sub2p by Hpr1p. Mol Cell Biol. 2002;22(23):8241–53. PubMed PMID: 12417727; PubMed Central PMCID: PMCPMC134069.

21. Lei EP, Krebber H, Silver PA. Messenger RNAs are recruited for nuclear export during transcription. Genes Dev. 2001;15(14):1771–82. doi: 10.1101/gad.892401. PubMed PMID: 11459827; PubMed Central PMCID: PMCPMC312744.

22. Huertas P, Aguilera A. Cotranscriptionally formed DNA:RNA hybrids mediate transcription elongation impairment and transcription-associated recombination. Mol Cell. 2003;12(3):711–21. PubMed PMID: 14527416.

23. Cerritelli SM, Crouch RJ. Ribonuclease H: the enzymes in eukaryotes. FEBS J. 2009;276(6):1494–505. doi: 10.1111/j.1742-4658.2009.06908.x. PubMed PMID: 19228196; PubMed Central PMCID: PMCPMC2746905.

24. Gomez-Gonzalez B, Garcia-Rubio M, Bermejo R, Gaillard H, Shirahige K, Marin A, et al. Genome-wide function of THO/TREX in active genes prevents R-loop-dependent replication obstacles. EMBO J. 2011;30(15):3106–19. doi: 10.1038/emboj.2011.206. PubMed PMID: 21701562; PubMed Central PMCID: PMCPMC3160181.

25. Wellinger RE, Prado F, Aguilera A. Replication fork progression is impaired by transcription in hyperrecombinant yeast cells lacking a functional THO complex. Mol Cell Biol. 2006;26(8):3327–34. doi: 10.1128/MCB.26.8.3327-3334.2006. PubMed PMID: 16581804; PubMed Central PMCID: PMCPMC1446968.

26. Luna R, Rondon AG, Perez-Calero C, Salas-Armenteros I, Aguilera A. The THO Complex as a Paradigm for the Prevention of Cotranscriptional R-Loops. Cold Spring Harb Symp Quant Biol. 2020. Epub 2020/06/05. doi: 10.1101/sqb.2019.84.039594. PubMed PMID: 32493765.

27. Pfeiffer V, Crittin J, Grolimund L, Lingner J. The THO complex component Thp2 counteracts telomeric R-loops and telomere shortening. EMBO J. 2013;32(21):2861–71. doi: 10.1038/emboj.2013.217. PubMed PMID: 24084588; PubMed Central PMCID: PMCPMC3817467.

28. Voynov V, Verstrepen KJ, Jansen A, Runner VM, Buratowski S, Fink GR. Genes with internal repeats require the THO complex for transcription. Proc Natl Acad Sci U S A. 2006;103(39):14423–8. Epub 2006/09/20. doi: 10.1073/pnas.0606546103. PubMed PMID: 16983072; PubMed Central PMCID: PMCPMC1599979.

29. Kilchert C, Wittmann S, Vasiljeva L. The regulation and functions of the nuclear RNA exosome complex. Nat Rev Mol Cell Biol. 2016;17(4):227–39. doi: 10.1038/nrm.2015.15. PubMed PMID: 26726035.

30. Schmidt K, Butler JS. Nuclear RNA surveillance: role of TRAMP in controlling exosome specificity. Wiley Interdiscip Rev RNA. 2013;4(2):217–31. doi: 10.1002/wrna.1155. PubMed PMID: 23417976; PubMed Central PMCID: PMCPMC3578152.

31. Delan-Forino C, Spanos C, Rappsilber J, Tollervey D. Substrate specificity of the TRAMP nuclear surveillance complexes. Nat Commun. 2020;11(1):3122. Epub 2020/06/21. doi: 10.1038/s41467-020-16965-4. PubMed PMID: 32561742; PubMed Central PMCID: PMCPMC7305330.

32. Kong KY, Tang HM, Pan K, Huang Z, Lee TH, Hinnebusch AG, et al. Cotranscriptional recruitment of yeast TRAMP complex to intronic sequences promotes optimal pre-mRNA splicing. Nucleic Acids Res. 2014;42(1):643–60. Epub 2013/10/08. doi: 10.1093/nar/gkt888. PubMed PMID: 24097436; PubMed Central PMCID: PMCPMC3874199.

33. Fox MJ, Mosley AL. Rrp6: Integrated roles in nuclear RNA metabolism and transcription termination. Wiley Interdiscip Rev RNA. 2016;7(1):91–104. doi: 10.1002/wrna.1317. PubMed PMID: 26612606; PubMed Central PMCID: PMCPMC4715707.

34. Houseley J, Kotovic K, El Hage A, Tollervey D. Trf4 targets ncRNAs from telomeric and rDNA spacer regions and functions in rDNA copy number control. EMBO J. 2007;26(24):4996–5006. Epub 2007/11/17. doi: 10.1038/sj.emboj.7601921. PubMed PMID: 18007593; PubMed Central PMCID: PMCPMC2080816.

35. Stuparevic I, Mosrin-Huaman C, Hervouet-Coste N, Remenaric M, Rahmouni AR. Cotranscriptional recruitment of RNA exosome cofactors Rrp47p and Mpp6p and two distinct Trf-Air-Mtr4 polyadenylation (TRAMP) complexes assists the exonuclease Rrp6p in the targeting and degradation of an aberrant messenger ribonucleoprotein particle (mRNP) in yeast. J Biol Chem. 2013;288(44):31816–29. Epub 2013/09/21. doi: 10.1074/jbc.M113.491290. PubMed PMID: 24047896; PubMed Central PMCID: PMCPMC3814775.

36. Gavalda S, Gallardo M, Luna R, Aguilera A. R-loop mediated transcription-associated recombination in trf4Delta mutants reveals new links between RNA surveillance and genome integrity. PLoS One. 2013;8(6):e65541. doi: 10.1371/journal.pone.0065541. PubMed PMID: 23762389; PubMed Central PMCID: PMCPMC3676323.

37. Wahba L, Amon JD, Koshland D, Vuica-Ross M. RNase H and multiple RNA biogenesis factors cooperate to prevent RNA:DNA hybrids from generating genome instability. Mol Cell. 2011;44(6):978–88. doi: 10.1016/j.molcel.2011.10.017. PubMed PMID: 22195970; PubMed Central PMCID: PMCPMC3271842.

38. Houseley J, LaCava J, Tollervey D. RNA-quality control by the exosome. Nat Rev Mol Cell Biol. 2006;7(7):529–39. doi: 10.1038/nrm1964. PubMed PMID: 16829983.

39. Luna R, Jimeno S, Marin M, Huertas P, Garcia-Rubio M, Aguilera A. Interdependence between transcription and mRNP processing and export, and its impact on genetic stability. Mol Cell. 2005;18(6):711–22. doi: 10.1016/j.molcel.2005.05.001. PubMed PMID: 15949445.

40. Wahba L, Gore SK, Koshland D. The homologous recombination machinery modulates the formation of RNA-DNA hybrids and associated chromosome instability. Elife. 2013;2:e00505. doi: 10.7554/eLife.00505. PubMed PMID: 23795288; PubMed Central PMCID: PMCPMC3679537.

41. Ogami K, Chen Y, Manley JL. RNA surveillance by the nuclear RNA exosome: mechanisms and significance. Noncoding RNA. 2018;4(1). Epub 2018/04/10. doi: 10.3390/ncrna4010008. PubMed PMID: 29629374; PubMed Central PMCID: PMCPMC5886371.

42. Larochelle M, Lemay JF, Bachand F. The THO complex cooperates with the nuclear RNA surveillance machinery to control small nucleolar RNA expression. Nucleic Acids Res. 2012;40(20):10240–53. doi: 10.1093/nar/gks838. PubMed PMID: 22965128; PubMed Central PMCID: PMCPMC3488260.

43. Callahan JL, Andrews KJ, Zakian VA, Freudenreich CH. Mutations in yeast replication proteins that increase CAG/CTG expansions also increase repeat fragility. Mol Cell Biol. 2003;23(21):7849–60. PubMed PMID: 14560028; PubMed Central PMCID: PMCPMC207578.

44. Sundararajan R, Gellon L, Zunder RM, Freudenreich CH. Double-strand break repair pathways protect against CAG/CTG repeat expansions, contractions and repeat-mediated chromosomal fragility in Saccharomyces cerevisiae. Genetics. 2010;184(1):65–77. doi: 10.1534/genetics.109.111039. PubMed PMID: 19901069; PubMed Central PMCID: PMCPMC2815931.

45. Gellon L, Razidlo DF, Gleeson O, Verra L, Schulz D, Lahue RS, et al. New functions of Ctf18-RFC in preserving genome stability outside its role in sister chromatid cohesion. PLoS Genet. 2011;7(2):e1001298. doi: 10.1371/journal.pgen.1001298. PubMed PMID: 21347277; PubMed Central PMCID: PMCPMC3037408.

46. Garcia-Rubio M, Chavez S, Huertas P, Tous C, Jimeno S, Luna R, et al. Different physiological relevance of yeast THO/TREX subunits in gene expression and genome integrity. Mol Genet Genomics. 2008;279(2):123–32. doi: 10.1007/s00438-007-0301-6. PubMed PMID: 17960421.

47. Tudek A, Lloret-Llinares M, Jensen TH. The multitasking polyA tail: nuclear RNA maturation, degradation and export. Philos Trans R Soc Lond B Biol Sci. 2018;373(1762). Epub 2018/11/07. doi: 10.1098/rstb.2018.0169. PubMed PMID: 30397105; PubMed Central PMCID: PMCPMC6232593.

48. Mitchell P. Exosome substrate targeting: the long and short of it. Biochem Soc Trans. 2014;42(4):1129–34. Epub 2014/08/12. doi: 10.1042/BST20140088. PubMed PMID: 25110014.

49. Polleys EJ, House NCM, Freudenreich CH. Role of recombination and replication fork restart in repeat instability. DNA Repair (Amst). 2017;56:156–65. doi: 10.1016/j.dnarep.2017.06.018. PubMed PMID: 28641941; PubMed Central PMCID: PMCPMC5541998.

50. Mosbach V, Poggi L, Richard GF. Trinucleotide repeat instability during double-strand break repair: from mechanisms to gene therapy. Curr Genet. 2019;65(1):17–28. Epub 2018/07/06. doi: 10.1007/s00294-018-0865-1. PubMed PMID: 29974202.

51. Gomez-Gonzalez B, Felipe-Abrio I, Aguilera A. The S-phase checkpoint is required to respond to R-loops accumulated in THO mutants. Mol Cell Biol. 2009;29(19):5203–13. doi: 10.1128/MCB.00402-09. PubMed PMID: 19651896; PubMed Central PMCID: PMCPMC2747986.

52. San Martin-Alonso M, Soler-Oliva ME, Garcia-Rubio M, Garcia-Muse T, Aguilera A. Harmful R-loops are prevented via different cell cycle-specific mechanisms. Nat Commun. 2021;12(1):4451. Epub 2021/07/24. doi: 10.1038/s41467-021-24737-x. PubMed PMID: 34294712; PubMed Central PMCID: PMCPMC8298424.

53. Janke C, Magiera MM, Rathfelder N, Taxis C, Reber S, Maekawa H, et al. A versatile toolbox for PCR-based tagging of yeast genes: new fluorescent proteins, more markers and promoter substitution cassettes. Yeast. 2004;21(11):947–62. doi: 10.1002/yea.1142. PubMed PMID: 15334558.

54. Gomez-Gonzalez B, Aguilera A. Activation-induced cytidine deaminase action is strongly stimulated by mutations of the THO complex. Proc Natl Acad Sci U S A. 2007;104(20):8409–14. doi: 10.1073/pnas.0702836104. PubMed PMID: 17488823; PubMed Central PMCID: PMCPMC1895963.

55. Castillo-Guzman D, Chedin F. Defining R-loop classes and their contributions to genome instability. DNA Repair (Amst). 2021;106:103182. Epub 2021/07/25. doi: 10.1016/j.dnarep.2021.103182. PubMed PMID: 34303066.

56. Karakasili E, Burkert-Kautzsch C, Kieser A, Strasser K. Degradation of DNA damage-independently stalled RNA polymerase II is independent of the E3 ligase Elc1. Nucleic Acids Res. 2014;42(16):10503–15. doi: 10.1093/nar/gku731. PubMed PMID: 25120264; PubMed Central PMCID: PMCPMC4176355.

57. Lalonde M, Trauner M, Werner M, Hamperl S. Consequences and Resolution of Transcription-Replication Conflicts. Life (Basel). 2021;11(7). Epub 2021/07/03. doi: 10.3390/life11070637. PubMed PMID: 34209204; PubMed Central PMCID: PMCPMC8303131.

58. Alam MS. Proximity Ligation Assay (PLA). Curr Protoc Immunol. 2018;123(1):e58. Epub 2018/09/22. doi: 10.1002/cpim.58. PubMed PMID: 30238640; PubMed Central PMCID: PMCPMC6205916.

59. Kotsantis P, Segura-Bayona S, Margalef P, Marzec P, Ruis P, Hewitt G, et al. RTEL1 Regulates G4/R-Loops to Avert Replication-Transcription Collisions. Cell Rep. 2020;33(12):108546. Epub 2020/12/29. doi: 10.1016/j.celrep.2020.108546. PubMed PMID: 33357438; PubMed Central PMCID: PMCPMC7773548.

60. Hamperl S, Bocek MJ, Saldivar JC, Swigut T, Cimprich KA. Transcription-Replication Conflict Orientation Modulates R-Loop Levels and Activates Distinct DNA Damage Responses. Cell. 2017;170(4):774–86 e19. Epub 2017/08/13. doi: 10.1016/j.cell.2017.07.043. PubMed PMID: 28802045; PubMed Central PMCID: PMCPMC5570545.

61. Manfrini N, Trovesi C, Wery M, Martina M, Cesena D, Descrimes M, et al. RNA-processing proteins regulate Mec1/ATR activation by promoting generation of RPA-coated ssDNA. EMBO Rep. 2015;16(2):221–31. doi: 10.15252/embr.201439458. PubMed PMID: 25527408; PubMed Central PMCID: PMCPMC4328749.

62. Domingo-Prim J, Endara-Coll M, Bonath F, Jimeno S, Prados-Carvajal R, Friedlander MR, et al. EXOSC10 is required for RPA assembly and controlled DNA end resection at DNA double-strand breaks. Nat Commun. 2019;10(1):2135. Epub 2019/05/16. doi: 10.1038/s41467-019-10153-9. PubMed PMID: 31086179; PubMed Central PMCID: PMCPMC6513946.

63. Deng SK, Chen H, Symington LS. Replication protein A prevents promiscuous annealing between short sequence homologies: Implications for genome integrity. Bioessays. 2015;37(3):305–13. Epub 2014/11/18. doi: 10.1002/bies.201400161. PubMed PMID: 25400143; PubMed Central PMCID: PMCPMC4410862.

64. Lin Y, Leng M, Wan M, Wilson JH. Convergent transcription through a long CAG tract destabilizes repeats and induces apoptosis. Mol Cell Biol. 2010;30(18):4435–51. doi: 10.1128/MCB.00332-10. PubMed PMID: 20647539; PubMed Central PMCID: PMCPMC2937530.

65. Khristich AN, Armenia JF, Matera RM, Kolchinski AA, Mirkin SM. Large-scale contractions of Friedreich’s ataxia GAA repeats in yeast occur during DNA replication due to their triplex-forming ability. Proc Natl Acad Sci U S A. 2020;117(3):1628–37. Epub 2020/01/09. doi: 10.1073/pnas.1913416117. PubMed PMID: 31911468; PubMed Central PMCID: PMCPMC6983365.

66. Branzei D, Szakal B. Building up and breaking down: mechanisms controlling recombination during replication. Crit Rev Biochem Mol Biol. 2017;52(4):381–94. Epub 2017/03/23. doi: 10.1080/10409238.2017.1304355. PubMed PMID: 28325102.

67. Bruning JG, Marians KJ. Replisome bypass of transcription complexes and R-loops. Nucleic Acids Res. 2020;48(18):10353–67. Epub 2020/09/15. doi: 10.1093/nar/gkaa741. PubMed PMID: 32926139; PubMed Central PMCID: PMCPMC7544221.

68. McGinty RJ, Puleo F, Aksenova AY, Hisey JA, Shishkin AA, Pearson EL, et al. A Defective mRNA Cleavage and Polyadenylation Complex Facilitates Expansions of Transcribed (GAA)n Repeats Associated with Friedreich’s Ataxia. Cell Rep. 2017;20(10):2490–500. doi: 10.1016/j.celrep.2017.08.051. PubMed PMID: 28877480; PubMed Central PMCID: PMCPMC5658003.

69. Gavalda S, Santos-Pereira JM, Garcia-Rubio ML, Luna R, Aguilera A. Excess of Yra1 RNA-Binding Factor Causes Transcription-Dependent Genome Instability, Replication Impairment and Telomere Shortening. PLoS Genet. 2016;12(4):e1005966. Epub 2016/04/02. doi: 10.1371/journal.pgen.1005966. PubMed PMID: 27035147; PubMed Central PMCID: PMCPMC4818039.

70. Garcia-Rubio M, Aguilera P, Lafuente-Barquero J, Ruiz JF, Simon MN, Geli V, et al. Yra1-bound RNA-DNA hybrids cause orientation-independent transcription-replication collisions and telomere instability. Genes Dev. 2018;32(13-14):965–77. Epub 2018/06/30. doi: 10.1101/gad.311274.117. PubMed PMID: 29954833; PubMed Central PMCID: PMCPMC6075034.

71. Katahira J, Senokuchi K, Hieda M. Human THO maintains the stability of repetitive DNA. Genes Cells. 2020;25(5):334–42. Epub 2020/02/18. doi: 10.1111/gtc.12760. PubMed PMID: 32065701.

72. Feretzaki M, Pospisilova M, Valador Fernandes R, Lunardi T, Krejci L, Lingner J. RAD51-dependent recruitment of TERRA lncRNA to telomeres through R-loops. Nature. 2020;587(7833):303–8. Epub 2020/10/16. doi: 10.1038/s41586-020-2815-6. PubMed PMID: 33057192.

73. Chappidi N, Nascakova Z, Boleslavska B, Zellweger R, Isik E, Andrs M, et al. Fork Cleavage-Religation Cycle and Active Transcription Mediate Replication Restart after Fork Stalling at Co-transcriptional R-Loops. Mol Cell. 2020;77(3):528–41 e8. Epub 2019/11/25. doi: 10.1016/j.molcel.2019.10.026. PubMed PMID: 31759821.

74. Mazina OM, Somarowthu S, Kadyrova LY, Baranovskiy AG, Tahirov TH, Kadyrov FA, et al. Replication protein A binds RNA and promotes R-loop formation. J Biol Chem. 2020;295(41):14203–13. Epub 2020/08/17. doi: 10.1074/jbc.RA120.013812. PubMed PMID: 32796030; PubMed Central PMCID: PMCPMC7549048.

75. Promonet A, Padioleau I, Liu Y, Sanz L, Biernacka A, Schmitz AL, et al. Topoisomerase 1 prevents replication stress at R-loop-enriched transcription termination sites. Nat Commun. 2020;11(1):3940. Epub 2020/08/10. doi: 10.1038/s41467-020-17858-2. PubMed PMID: 32769985; PubMed Central PMCID: PMCPMC7414224.

76. Kemiha S, Poli J, Lin YL, Lengronne A, Pasero P. Toxic R-loops: Cause or consequence of replication stress? DNA Repair (Amst). 2021;107:103199. Epub 2021/08/17. doi: 10.1016/j.dnarep.2021.103199. PubMed PMID: 34399314.

77. Longtine MS, McKenzie A, 3rd, Demarini DJ, Shah NG, Wach A, Brachat A, et al. Additional modules for versatile and economical PCR-based gene deletion and modification in Saccharomyces cerevisiae. Yeast. 1998;14(10):953–61. doi: 10.1002/(SICI)1097-0061(199807)14:10<953::AID-YEA293>3.0.CO;2-U. PubMed PMID: 9717241.

78. Polleys EJ, Freudenreich CH. Methods to Study Repeat Fragility and Instability in Saccharomyces cerevisiae. Methods Mol Biol. 2018;1672:403–19. doi: 10.1007/978-1-4939-7306-4_28. PubMed PMID: 29043639.

79. Polleys EJ, Freudenreich CH. Genetic Assays to Study Repeat Fragility in Saccharomyces cerevisiae. In: Richard G-F, editor.Trinucleotide Repeats: Methods and Protocols. New York, NY: Springer New York; 2020. p. 83–101.

80. Hall BM, Ma CX, Liang P, Singh KK. Fluctuation analysis CalculatOR: a web tool for the determination of mutation rate using Luria-Delbruck fluctuation analysis. Bioinformatics. 2009;25(12):1564–5. Epub 2009/04/17. doi: 10.1093/bioinformatics/btp253. PubMed PMID: 19369502; PubMed Central PMCID: PMCPMC2687991.

81. Radchenko EA, McGinty RJ, Aksenova AY, Neil AJ, Mirkin SM. Quantitative Analysis of the Rates for Repeat-Mediated Genome Instability in a Yeast Experimental System. In: Muzi-Falconi M, Brown GW, editors. Genome Instability: Methods and Protocols. New York, NY: Springer New York; 2018. p. 421–38.

82. Boguslawski SJ, Smith DE, Michalak MA, Mickelson KE, Yehle CO, Patterson WL, et al. Characterization of monoclonal antibody to DNA.RNA and its application to immunodetection of hybrids. J Immunol Methods. 1986;89(1):123–30. PubMed PMID: 2422282.

83. Hu Z, Zhang A, Storz G, Gottesman S, Leppla SH. An antibody-based microarray assay for small RNA detection. Nucleic Acids Res. 2006;34(7):e52. doi: 10.1093/nar/gkl142. PubMed PMID: 16614443; PubMed Central PMCID: PMCPMC1435976.

84. Alberts N, Mathangasinghe Y, Nillegoda NB. In Situ Monitoring of Transiently Formed Molecular Chaperone Assemblies in Bacteria, Yeast, and Human Cells. J Vis Exp. 2019;(151). Epub 2019/09/17. doi: 10.3791/60172. PubMed PMID: 31524873.

