## Supplemental Information for "THO and TRAMP complexes prevent transcription-replication conflicts, DNA breaks, and CAG repeat contractions"

### Supplementary Tables

**Table S1.** Fragility analysis of CAG-70 repeats on the *URA3*-YAC.

| **Strains** | **Individual rate of FOA$^R$ (x10$^{-6}$)** | **Average rate of FOA$^R$ (x10$^{-6}$) ± S.E.M** | **Fold over wt** | **# of Assays** | **p-value to wt** | **Other p-values a to *rnh1Δrnh201Δ;* b to *thp2Δ*; c to *trf4Δ*** |
|---|---|---|---|---|---|---|
| Wild-type | 4.2[1] | 7.01±1.3 | - | 5 | - | - |
| | 5.2[1] | | | | | |
| | 8.5[1] | | | | | |
| | 5.97 | | | | | |
| | 11.2 | | | | | |
| *rnh1Δrnh201Δ*[2] | 57.6 | 48.7±6.9 | 6.9 | 3 | $2.3x10^{-4}$ | - |
| | 35.1 | | | | | |
| | 53.4 | | | | | |
| *thp2Δ* | 33.4 | 33.2±3.5 | 4.7 | 6 | $1.1x10^{-4}$ | 0.057[a] |
| | 40.4 | | | | | |
| | 44.8 | | | | | |
| | 31.3 | | | | | |
| | 21.1 | | | | | |
| | 28.1 | | | | | |
| *mft1Δ* | 43.3 | 47.4±3.9 | 6.8 | 4 | $1.2x10^{-5}$ | 0.87[a]; 0.028[b] |
| | 52.3 | | | | | |
| | 55.3 | | | | | |
| | 38.7 | | | | | |
| *thp2Δrnh1Δrnh201Δ* | 475 | 282±71 | 40.0 | 4 | $3.1x10^{-3}$ | 0.039[a]; $2.2X10^{-3}$[b] |
| | 187 | | | | | |
| | 167 | | | | | |
| | 297 | | | | | |
| *trf4Δ* | 39.1 | 52.8±9.8 | 7.5 | 4 | $1.2x10^{-3}$ | 0.77[a]; 0.059[b] |
| | 62.7 | | | | | |
| | 75.3 | | | | | |
| | 33.9 | | | | | |
| *trf5Δ* | 12.9 | 12.4±1.8 | 1.8 | 3 | 0.045 | 0.018[c] |
| | 9.1 | | | | | |
| | 15.2 | | | | | |
| *rrp6Δ* | 56.5 | 56.6±0.2 | 8.1 | 3 | $1.1x10^{-7}$ | 0.76[c] |
| | 57 | | | | | |
| | 56.2 | | | | | |
| *trf4Δrnh1Δrnh201Δ* | 340 | 552±74 | 74.4 | 4 | $9.9x10^{-5}$ | $3.0X10^{-3}$[a]; $8.0X10^{-4}$[c] |
| | 552 | | | | | |
| | 699 | | | | | |
| | 498 | | | | | |
| *trf4Δrrp6Δ* | 113 | 136±28 | 19.3 | 4 | $1.2x10^{-3}$ | 0.030[c]; 0.061 to *rrp6Δ* |
| | 84 | | | | | |
| | 133 | | | | | |

| | | | | | | |
|---|---|---|---|---|---|---|
| | 214 | | | | | |
| *rad51Δ* | 14.9 | 21.2±4.5 | 3.0 | 3 | 8.3x10$^{-3}$ | - |
| | 29.9 | | | | | |
| | 18.9 | | | | | |
| *thp2Δrad51Δ* | 55.0 | 40.9±3.6 | 5.8 | 5 | 2.2x10$^{-5}$ | 0.16[b]; 0.015 to *rad51Δ* |
| | 39.1 | | | | | |
| | 33.9 | | | | | |
| | 37.3 | | | | | |
| | 39.4 | | | | | |
| *trf4Δrad51Δ* | 29.5 | 28.1±4.3 | 4.0 | 4 | 1.2x10$^{-3}$ | 0.06[c]; 0.32 to *rad51Δ* |
| | 38.9 | | | | | |
| | 25.5 | | | | | |
| | 18.5 | | | | | |
| **Strains used for one-colony fragility assays** | | | | | | **c to *thp2Δ*; d to *trf4Δ*** |
| *thp2Δ* | 0.0435 | 0.199±0.10 | - | 5 | - | - |
| | 0.117 | | | | | |
| | 0.594 | | | | | |
| | 0.149 | | | | | |
| | 0.092 | | | | | |
| *trf4Δ* | 0.232 | 0.319±0.031 | - | 4 | - | - |
| | 0.319 | | | | | |
| | 0.363 | | | | | |
| | 0.361 | | | | | |
| *thp2Δtrf4Δ* | 1900 | 2300±350 | - | 3 | - | 1.1x10$^{-4}$[c]; 5.5x10$^{-4}$[d] |
| | 3000 | | | | | |
| | 2000 | | | | | |
| **Strains with p*MET25-RNH1*** | | | | | | **to +Met condition in the same mutant** |
| wild-type in +Met | 18.2 | 24.9±3.7 | - | 4 | - | - |
| | 30.9 | | | | | |
| | 25.7 | | | | | |
| wild-type in -Met | 16.2 | 24.9±4.7 | - | 4 | - | 0.99 |
| | 32.5 | | | | | |
| | 30.9 | | | | | |
| *thp2Δ* in +Met | 36.3 | 32.6±2.9 | - | 3 | - | - |
| | 26.8 | | | | | |
| | 34.7 | | | | | |
| *thp2Δ* in -Met | 14.0 | 19.2±2.6 | - | 3 | - | 0.027 |
| | 20.8 | | | | | |
| | 22.7 | | | | | |
| *trf4Δ* in +Met | 55.0 | 54.6±9.4 | - | 3 | - | - |
| | 38.1 | | | | | |
| | 70.6 | | | | | |
| *trf4Δ* in -Met | 45.6 | 48.3±4.0 | - | 3 | - | 0.57 |

| | | | | | | |
|---|---|---|---|---|---|---|
| | 43.1 | | | | | |
| | 56.1 | | | | | |
| **Strains with RPA overexpression vector** | | | | | | **to no overexpression condition** |
| Wild-type RPA overexpressed | 24.3 | 11.2±4.5 | - | 4 | - | 0.36 |
| | 9.72 | | | | | |
| | 5.95 | | | | | |
| | 4.69 | | | | | |
| *trf4Δ* RPA overexpressed | 34.9 | 23.2±4.5 | - | 3 | - | 0.034 |
| | 23.8 | | | | | |
| | 13.0 | | | | | |
| | 20.9 | | | | | |

1, indicated wild-type data are from (Kerrest et al. 2009); 2, data from (Su and Freudenreich 2017); "-Met" represents yeast synthetic media lacking methionine, which confers a *RNH1* over-expression condition; "+Met" represents yeast synthetic media containing methionine, thus does not induce *RNH1* over-expression (see Fig. S3).

**Table S2.** Fragility analysis of no tract (CAG-0) on the *URA3*-YAC.

| **Strains** | Individual rate of $FOA^R$ $(X10^{-6})$ | Average rate of $FOA^R$ $(X10^{-6})$ ± S.E.M | Fold over wt | # of Assays | p-value to wt | Other p-values c, to *trf4Δ* |
|---|---|---|---|---|---|---|
| Wild-type[1] | 2.6 | 2.2±0.45 | - | 3 | - | - |
| | 2.8 | | | | | |
| | 1.3 | | | | | |
| *thp2Δ* | 5.5 | 9.1±0.98 | 4.1 | 7 | $2.4X10^{-3}$ | - |
| | 8.6 | | | | | |
| | 11.7 | | | | | |
| | 10.8 | | | | | |
| | 12.5 | | | | | |
| | 6.8 | | | | | |
| | 8.1 | | | | | |
| *trf4Δ* | 7.0 | 10.8±1.8 | 4.9 | 3 | 0.020 | - |
| | 14.7 | | | | | |
| | 10.6 | | | | | |

1, wild-type data are from (Kerrest et al. 2009).

**Table S3.** Instability analysis of CAG-70 repeats on the *URA3*-YAC.

| | | **Contractions** | | | | **Expansions** | | | |
|---|---|---|---|---|---|---|---|---|---|
| **Strain** | Total rxns | # | % | Fold over wt | p-value to wt | # | % | Fold over wt | p-value to wt |
| wild-type[1] | 460 | 20 | 4.3 | - | - | 5 | 1.1 | - | - |
| *thp2Δ* | 256 | 29 | 11.3 | 2.6 | $6.0X10^{-3}$ | 8 | 3.1 | 2.8 | 0.076 |
| *trf4Δ* | 130 | 20 | 15.4 | 3.6 | $1.0X10^{-3}$ | 4 | 3.1 | 2.8 | 0.11 |

1, wild-type data are from (Kerrest et al. 2009)

**Table S4.** Analysis of *URA3* presence in FOA-resistant colonies.

| **Strain** | Presence of *URA3* | | Total number of FOA$^R$ colonies[1] | Percent end loss (no *URA3*) | Method used |
|---|---|---|---|---|---|
| | number | percent | | | |
| Wild-type | 0 | 0% | 20 | 100% | PCR |
| *rnh1Δrnh201Δ* | 0 | 0% | 20 | 100% | PCR |
| *thp2Δ* | 0 | 0% | 10 | 100% | Southern Blot |
| *trf4Δ* | 0 | 0% | 30 | 100% | PCR |
| Wild-type RPA overexpressed | 3 | 7.5% | 40 | 92.5% | PCR |
| *trf4Δ* RPA overexpressed | 6 | 15% | 40 | 85% | PCR |

1, Only one FOA$^R$ colony per plate was tested, assuring that each event was independent (For 10-colony assays, each culture plated on each FOA-Leu plate is from an individual parent colony). PCR methods and primers used for checking *URA3* locus are listed in Supplementary Table 4. Southern blot method is the same as in (Callahan et al. 2003).

**Table S5.** Viability of yeast strains on YC-Leu.

| Strain | % Viability | Average % Viability |
|---|---|---|
| wild-type | 52.6 | 76.5 |
| | 107 | |
| | 70.0 | |
| *thp2ΔrnhΔ* | 22.5 | 27.2 |
| | 32.0 | |
| | 27.0 | |
| *trf4ΔrnhΔ* | 20.0 | 22.0 |
| | 13.0 | |
| | 33.0 | |
| *thp2Δtrf4Δ* | 13.0 | 10.0 |
| | 10.0 | |
| | 7.00 | |

**Table S6.** DNA: RNA immunoprecipitation (DRIP) analysis.

| Locus | Strain | Individual signal (over *MMR1*) | Average signal (over *MMR1*) ± S.E.M | p-value compared to wild-type |
|---|---|---|---|---|
| Cross G4T4 | wild-type | 0.55 | 0.74±0.19 | - |
| | | 0.92 | | |
| | *thp2Δ* | 1.74 | 2.05±0.30 | 0.07 |
| | | 2.35 | | |
| | *trf4Δ* | 2.34 | 1.09±0.50 | 0.67 |
| | | 1.44 | | |
| | | 0.29 | | |
| | | 0.28 | | |
| *URA3* locus | wild-type | 1.50 | 1.63±0.13 | - |
| | | 1.75 | | |
| | *thp2Δ* | 1.51 | 2.20±0.69 | 0.50 |

| | | | | |
|---|---|---|---|---|
| | | 2.88 | | |
| | *trf4Δ* | 1.47 | 0.86±0.17 | 0.05 |
| | | 0.99 | | |
| | | 0.64 | | |
| | | 0.65 | | |
| | | 0.54 | | |
| CAG-70 locus | wild-type | 2.67 | 2.56±0.16 | - |
| | | 2.24 | | |
| | | 2.78 | | |
| | *thp2Δ* | 3.08 | 2.98±0.10 | 0.16 |
| | | 2.88 | | |
| | *trf4Δ* | 6.89 | 2.80±1.17 | 0.88 |
| | | 0.94 | | |
| | | 0.86 | | |
| | | 1.32 | | |
| | | 3.99 | | |
| | *rnh1Δrnh201Δ** | 9.24 | - | - |

*Other *rnh1Δrnh201Δ* data points found in (Su and Freudenreich 2017). Data point shown in table for *rnh1Δrnh201Δ* was collected alongside wild-type, *thp2Δ*, and *trf4Δ* as a control.

**Table S7.** *RNH1* expression data.

| Strain | % ACT1 (by Absolute Quantity) | % ACT1 Average | Fold over no RNH1 induction (+Met condition) |
|---|---|---|---|
| wild-type (+Met) | 9.70 | 9.29 | - |
| | 8.88 | | |
| wild-type (-Met) | 28.6 | 30.1 | 3.24 |
| | 31.5 | | |
| *thp2Δ* (+Met) | 9.75 | 6.74 | - |
| | 3.72 | | |
| *thp2Δ* (-Met) | 31.7 | 32.8 | 4.87 |
| | 33.9 | | |
| *trf4Δ* (+Met) | 12.4 | 9.1 | - |
| | 5.75 | | |
| *trf4Δ* (-Met) | 30.6 | 31.0 | 3.41 |
| | 31.3 | | |

**Table S8.** Chromatin immunoprecipitation (ChIP) analysis at CAG locus.

| ChIP protein | Strain | Individual signal (over ACT1) | Average signal (over ACT1) ± S.E.M | p-value compared to wild-type |
|---|---|---|---|---|
| RPA | wild-type | 1.10 | 1.25±0.10 | - |
| | | 1.50 | | |
| | | 1.30 | | |
| | | 1.10 | | |
| | *trf4Δ* | 0.89 | 0.91±0.03 | 0.03 |
| | | 0.97 | | |

| | | | | |
|---|---|---|---|---|
| | | 0.87 | | |
| | *rrp6Δ* | 1.06 | 1.08±0.02 | 0.18 |
| | | 1.09 | | |
| | *trf5Δ* | 1.16 | 1.10±0.06 | 0.37 |
| | | 1.04 | | |
| | *thp2Δ* | 1.37 | 1.30±0.08 | 0.78 |
| | | 1.22 | | |
| | *trf4Δthp2Δ* | 0.84 | 0.87±0.03 | 0.05 |
| | | 0.90 | | |
| RNAPII | wild-type | 1.04 | 1.41±0.40 | - |
| | | 2.21 | | |
| | | 0.99 | | |
| | *trf4Δ* | 3.82 | 3.89±0.49 | 0.017 |
| | | 4.77 | | |
| | | 3.08 | | |
| | *thp2Δ* | 2.81 | 4.11±1.18 | 0.05 |
| | | 6.47 | | |
| | | 3.04 | | |
| | *rnh1Δrnh201Δ* | 1.98 | 1.73±0.23 | 0.53 |
| | | 1.94 | | |
| | | 1.27 | | |

**Table S9.** Proximity-Ligation Assay (PLA) data.

| **Strains** | Antibody | Total # of cells | Total # of foci | Average foci per nucleus | Std Dev | p-value compared to wild-type | p-value compared to no *RNH1* or RPA over-expression | p-value compared to double antibody condition |
|---|---|---|---|---|---|---|---|---|
| wild-type | Both | 300 | 559 | 1.86 | 1.7 | - | - | - |
| | PCNA only | 300 | 6 | 0.0200 | 0.14 | - | - | <0.0001 |
| | RNAPII only | 300 | 20 | 0.0667 | 0.29 | - | - | <0.0001 |
| *thp2Δ* | Both | 300 | 890 | 2.97 | 2.0 | <0.0001 | - | - |
| | PCNA only | 300 | 44 | 0.147 | 0.50 | - | - | <0.0001 |
| | RNAPII only | 300 | 50 | 0.167 | 0.44 | - | - | <0.0001 |
| *trf4Δ* | Both | 400 | 1682 | 4.21 | 2.7 | <0.0001 | - | - |
| | PCNA only | 300 | 21 | 0.0700 | 0.31 | - | - | <0.0001 |
| | RNAPII only | 300 | 14 | 0.0467 | 0.23 | - | - | <0.0001 |
| *rnh1Δrnh201Δ* | Both | 300 | 730 | 2.43 | 1.9 | 0.0001 | - | - |
| | PCNA only | 300 | 0 | 0 | 0 | - | - | <0.0001 |

| | | | | | | | | |
|---|---|---|---|---|---|---|---|---|
| | RNAPII only | 300 | 8 | 0.0267 | 0.16 | - | - | <0.0001 |
| wild-type *pMET25-RNH1* | Both | 300 | 484 | 1.61 | 1.5 | 0.11 | 0.11 | - |
| | PCNA only | 300 | 3 | 0.0100 | 0.10 | - | - | <0.0001 |
| | RNAPII only | 300 | 5 | 0.0167 | 0.13 | - | - | <0.0001 |
| *thp2Δ pMET25-RNH1* | Both | 300 | 442 | 1.47 | 1.5 | 0.0038 | <0.0001 | - |
| | PCNA only | 300 | 1 | 0.00333 | 0.058 | - | - | <0.0001 |
| | RNAPII only | 300 | 1 | 0.00333 | 0.058 | - | - | <0.0001 |
| *trf4Δ pMET25-RNH1* | Both | 300 | 571 | 1.90 | 1.8 | 0.89 | <0.0001 | - |
| | PCNA only | 300 | 3 | 0.0100 | 0.10 | - | - | <0.0001 |
| | RNAPII only | 300 | 8 | 0.0267 | 0.16 | - | - | <0.0001 |
| wild-type RPA overexpressed | Both | 300 | 420 | 1.40 | 1.4 | 0.0011 | 0.0011 | - |
| | PCNA only | 300 | 6 | 0.0200 | 0.16 | - | - | <0.0001 |
| | RNAPII only | 300 | 1 | 0.00333 | 0.058 | - | - | <0.0001 |
| *trf4Δ* RPA overexpressed | Both | 300 | 686 | 2.29 | 2.0 | 0.014 | <0.0001 | - |
| | PCNA only | 300 | 8 | 0.0267 | 0.16 | - | - | <0.0001 |
| | RNAPII only | 300 | 3 | 0.0100 | 0.10 | - | - | <0.0001 |

**Table S10.** Quantification of foci detected outside of nucleus in PLA experiments.

| **Strains** | Average # foci in nucleus | Average # foci outside nucleus | Total # foci outside of nucleus | Total # of foci inside nucleus | Ratio of foci outside vs. inside nucleus |
|---|---|---|---|---|---|
| wild-type | 1.9 | 0.55 | 164 | 559 | 0.29 |
| *rnh1Δrnh201Δ* | 2.4 | 0.46 | 139 | 730 | 0.21 |
| *thp2Δ* | 3.0 | 0.61 | 184 | 890 | 0.28 |
| *trf4Δ* | 4.2 | 1.2 | 465 | 1682 | 0.19 |

Foci outside of the nucleus were not counted in the total # of foci presented in Table S9. We quantified these foci to ensure the percentage of foci outside of the nucleus did not vary significantly between strains.

**Table S11.** RPA expression data.

| Strain | % *ACT1* (by Absolute Quantity) | % *ACT1* Average | Fold Change over wild-type (by Absolute Quantity) | Fold change over wild-type Average |
|---|---|---|---|---|

| | | | | |
|---|---|---|---|---|
| *RFA1* | | | | |
| wild-type +RPA plasmid | 26.8 | 29.1 | 20.0 | 26.6 |
| | 9.85 | | 193 | |
| | 50.6 | | 26.7 | |
| *trf4Δ* +RPA plasmid | 37.1 | 44.5 | 27.6 | 40.7 |
| | 13.3 | | 261 | |
| | 83.1 | | 43.9 | |
| wild-type (no plasmid) | 1.34 | 1.09 | 1 | - |
| | 0.0511 | | 1 | |
| | 1.89 | | 1 | |
| *RFA2* | | | | |
| wild-type +RPA plasmid | 52.4 | 52.0 | 22.7 | 25.8 |
| | 27.4 | | 155 | |
| | 76.0 | | 21.3 | |
| *trf4Δ* +RPA plasmid | 46.0 | 81.0 | 19.9 | 40.1 |
| | 40.9 | | 230 | |
| | 156.0 | | 43.7 | |
| wild-type (no plasmid) | 2.31 | 2.02 | 1 | - |
| | 0.177 | | 1 | |
| | 3.57 | | 1 | |
| *RFA3* | | | | |
| wild-type +RPA plasmid | 37.2 | 43.7 | 12.5 | 19.4 |
| | 13.1 | | 90.6 | |
| | 80.7 | | 22.2 | |
| *trf4Δ* +RPA plasmid | 33.4 | 53.7 | 11.2 | 23.8 |
| | 15.5 | | 107 | |
| | 112 | | 30.9 | |
| wild-type (no plasmid) | 2.98 | 2.25 | 1 | - |
| | 0.145 | | 1 | |
| | 3.63 | | 1 | |

**Table S12.** Primers used in this study.

| Locus | Primer name | Oligonucleotide sequence |
|---|---|---|
| Cross CAG (instability) | NewCAGfor | CCTCAGCCTGGCCGAAAGAAAGAAA |
| | NewCAGrev | CAGTCACGACGTTGTAAAACGACGG |
| Cross CAG (confirming tract length for fragility assays) | CAG-2Step-F | GCGTGGAGGATGGAACACGGACGG |
| | CAG-2Step-R | GAAAGGGGGATGTGCTGCAAGGCG |
| Cross CAG (qPCR) | T7-20B | GAATTCGAGCTCCACCGCGG |
| | CTG rev2 | CCCAGGCCTCCAGTTTGC |
| Cross G4T4 (qPCR) | G4T4 right 65bp | CCTGTCGTGCCAGTGTATAC |
| | G4T4 left 90bp | GTGGCCAGGACCCAACGCTG |
| *MMR1* internal (qPCR) | MMR1 internal for | GCCCTAAGACTAGACTGGCAC |
| | MMR1 internal rev | GCAGAAGTTGGCTCCTCTTC |

| | | |
|---|---|---|
| *ACT1* internal (qPCR) | ACT1for3 | TCCAGATGGTCAAGTCATCA |
| | ACT1rev3 | TCGGCAATACCTGGGAACAT |
| *URA3* internal (check presence) | URA3 for2 | TGCTGCTACTCATCCTAG |
| | URA3 rev | TCCCAGCCTGCTTTTCTGTA |
| *RNH1* internal (qPCR) | RNH1-upreg-verif-Reverse | GCTTGTAATCATGCGCACTCATAC |
| | RNH1 internal forward 1 | GCGAGTTCATCGAAGGAATCGGC |

**Table S13.** Yeast Strains used in this study.

| Strain Number | Strain Background | Genotype | Reference |
|---|---|---|---|
| CFY765 | BY4705 | MATα, *ade2Δ::hisG, his3Δ200*; *leu2Δ0, lys2Δ0, met15Δ0, trp1Δ63, ura3Δ0, can*$^{R}$; YAC: *ade3-2p, LEU2,* CAG-0, *URA3*. | (Kerrest et al. 2009; Sundararajan et al. 2010) |
| CFY766 | BY4705 | MATα, *ade2Δ::hisG, his3Δ200*; *leu2Δ0, lys2Δ0, met15Δ0, trp1Δ63, ura3Δ0, can*$^{R}$; YAC: *ade3-2p, LEU2,* CAG-70,*URA3*. | (Kerrest et al. 2009; Sundararajan et al. 2010) |
| CFY3418, 3419 | BY4705 | CFY766, *rnh1Δ::His3MX6*; *rnh201Δ::TRP1* | (Su and Freudenreich 2017) |
| CFY2393, 2394 | BY4705 | CFY765, *thp2Δ::KanMX6* | This Study |
| CFY2395, 2396 | BY4705 | CFY766, *thp2Δ::KanMX6* | This Study |
| CFY4094, 4095 | BY4705 | CFY766, *mft1Δ::KanMX6* | This Study |
| CFY3416, 3417 | BY4705 | CFY3418, *thp2Δ::KanMX6* | This Study |
| CFY1976, 1977 | BY4705 | CFY765, *trf4Δ::His3MX6* | This Study |
| CFY1863 | BY4705 | CFY766, *trf4Δ::KanMX6* | This Study |
| CFY4113,4127 | BY4705 | CFY3418 *trf4Δ::KanMX6* | This Study |
| CFY4111, 4112 | BY4705 | CFY2395 *trf4Δ::His3MX6* | This Study |
| CFY2044, 2045 | BY4705 | CFY766 *trf5Δ::His3MX6* | This Study |
| CFY4224, 4225 | BY4705 | CFY766 *rrp6Δ::TRP1* | This Study |
| CFY4324, 4325 | BY4705 | CFY1977 *rrp6Δ::TRP1* | This Study |
| CFY3466 | BY4705 | MATα, *ade2Δ::hisG, his3Δ200*; leu*2Δ0, lys2Δ0, trp1Δ63, ura3Δ0, can*$^{R}$; YAC: *ade3-2p, LEU2,* CAG-70,*URA3*; *pMET25-RNH1*; *thp2Δ::KanMX6* | This Study |
| CFY4343, 4344 | BY4705 | MATα, *ade2Δ::hisG, his3Δ200*; *leu2Δ0, lys2Δ0, trp1Δ63, ura3Δ0, can*$^{R}$; YAC: *ade3-2p, LEU2,* CAG-70,*URA3*; *pMET25-RNH1*; *trf4Δ:: His3MX6* | This Study |

| | | | |
|---|---|---|---|
| CFY4334 | BY4705 | MATα, *ade2Δ::hisG, his3Δ200*; *leu2Δ0, lys2Δ0, trp1Δ63, ura3Δ0, can*$^{R}$; YAC: *ade3-2p, LEU2,* CAG-70,*URA3*; *pMET25-RNH1* | This Study |
| - | BY4705 | CFY766, transformed with pUC57 RPA OE *HIS3* plasmid containing *RFA1, RFA2, RFA3* genes (pUC57 made by S. Khristich, plasmid stock CFP772) | This Study, (Khristich et al. 2020)* |
| - | BY4705 | CFY1863, transformed with pUC57 RPA OE *HIS3* plasmid containing *RFA1, RFA2, RFA3* genes | This Study, (Khristich et al. 2020)* |

*For RPA overexpression experiments transformations were performed prior to each experiment with plasmid and strains that are listed.

**References**

Callahan JL, Andrews KJ, Zakian VA, Freudenreich CH. 2003. Mutations in yeast replication proteins that increase CAG/CTG expansions also increase repeat fragility. *Mol Cell Biol* **23**: 7849-7860.

Kerrest A, Anand RP, Sundararajan R, Bermejo R, Liberi G, Dujon B, Freudenreich CH, Richard GF. 2009. SRS2 and SGS1 prevent chromosomal breaks and stabilize triplet repeats by restraining recombination. *Nat Struct Mol Biol* **16**: 159-167.

Khristich AN, Armenia JF, Matera RM, Kolchinski AA, Mirkin SM. 2020. Large-scale contractions of Friedreich's ataxia GAA repeats in yeast occur during DNA replication due to their triplex-forming ability. *Proc Natl Acad Sci U S A* **117**: 1628-1637.

Su XA, Freudenreich CH. 2017. Cytosine deamination and base excision repair cause R-loop-induced CAG repeat fragility and instability in Saccharomyces cerevisiae. *Proc Natl Acad Sci U S A* **114**: E8392-E8401.

Sundararajan R, Gellon L, Zunder RM, Freudenreich CH. 2010. Double-strand break repair pathways protect against CAG/CTG repeat expansions, contractions and repeat-mediated chromosomal fragility in Saccharomyces cerevisiae. *Genetics* **184**: 65-77.
